# Enhancer status in the primitive endoderm supports unrestricted lineage plasticity in regulative development

**DOI:** 10.1101/2023.05.20.540779

**Authors:** Madeleine Linneberg-Agerholm, Annika Charlotte Sell, Alba Redo-Riveiro, Martin Proks, Teresa E. Knudsen, Marta Perera, Joshua M. Brickman

## Abstract

Mammalian blastocyst formation involves the specification of trophectoderm followed by the differentiation of the inner cell mass into either epiblast or primitive endoderm. During this time, the embryo maintains a window of plasticity and can redirect its cellular fate when challenged experimentally. In this context, we found that the primitive endoderm alone was sufficient to regenerate a complete blastocyst and continue normal postimplantation development to term. We identify an *in vitro* population similar to the early primitive endoderm *in vivo*, that exhibits the same embryonic and extra-embryonic potency, forming three dimensional embryoid structures. Commitment in early primitive endoderm is suppressed by JAK/STAT signalling, collaborating with OCT4 to safeguard enhancer status enabling multi-lineage differentiation. Our observations support the notion that transcription factor persistence underlies plasticity in regulative development and highlights the importance of primitive endoderm in perturbed development.

## Introduction

The early murine embryo possesses the remarkable ability to adapt and modulate its fate following various perturbations, and to compensate for loss of specific cell types. This feature is referred to as plasticity and enables cells to change their differentiation trajectory, a hallmark of regulative development. At the pinnacle of the cell potency hierarchy sits the 2 cell stage blastomere, which has the capacity to generate a complete organism from a single cell^1, 2^. The potential of each blastomere is then progressively restricted with every cell division and subsequent lineage segregation of the preimplantation embryo^3–7^.

The first lineage specification event takes place after morula compaction and the acquisition of polarity, leading to the formation of extra-embryonic trophectoderm (TE) and the inner cell mass (ICM). This is followed by blastulation, where the blastocoel forms alongside FGF/ERK-mediated differentiation of ICM cells to pluripotent epiblast (Epi), that gives rise to the majority of the embryo proper, or the bipotent extra-embryonic primitive endoderm (PrE), that later forms the parietal endoderm (PE) and visceral endoderm (VE) (reviewed by Chazaud and Yamanaka, 2016). During this process, cells of the Epi and PrE maintain a window of plasticity and differentiation potential when challenged experimentally, where both cell types exhibit multilineage potency upon chimera formation^9–11^. In unperturbed development, PrE-to-Epi switching is observed, but never vice versa, and only until the mid-blastocyst stage^12, 13^. The plasticity of PrE and Epi cells is progressively restricted throughout blastocyst maturation, and by the late blastocyst stage, both lineages are committed to their respective fates.

Oct4/Pou5f1 is a homeodomain transcription factor known for its role in supporting pluripotency *in vivo* and in embryonic stem cells (ESCs), as well as supporting transcription factor-mediated reprogramming of somatic cells to induced pluripotent stem cells (iPSCs)^14–17^. Zygotic Oct4 expression is activated prior to the 8-cell stage and maintained in all cells of the embryo^18^. OCT4 is initially expressed in both the early Epi and PrE cells^9, 18–20^, where it is required cell-autonomously during PrE specification to transduce FGF/ERK signalling in the nascent PrE^19^. OCT4 is also required for PrE induction *in vitro*, where alternative partnering of OCT4 with SOX2 and SOX17 allows the expression of endodermal genes^21^. Following PrE specification, OCT4 becomes restricted to the Epi compartment, coinciding with observed loss of plasticity during blastocyst maturation^9^. While the role of Oct4 in pluripotency and reprogramming has been extensively studied, the meaning of its expression in the early endoderm remains an enigma.

Naïve ESCs are immortal cell lines derived from the ICM that closely resemble the peri implantation Epi and represent an important model for early development used extensively for further *in vitro* differentiation^22^. These can be cultured in a variety of conditions^23^, including the so-called ground state, in which ESCs are maintained in defined conditions with the cytokine LIF and inhibitors of GSK3 and MEK (2iLIF)^24^. Trophoblast stem cells (TSCs) recapitulating the late TE and early extra-embryonic ectoderm^25^, and extra embryonic endoderm (XEN) cells that resemble the postimplantation-stage PE^26, 27^ have similarly been reported. We recently described PrE stem cells that can be derived from naïve ESCs and expanded as blastocyst-stage PrE, termed naïve extra-embryonic endoderm (nEnd), through LIF, Wnt and TGFβ signalling^28^. Where ESCs are known to exhibit heterogeneities that reflect differentiation competence, it is likely also a general property of stem cell models including nEnd.

In this paper, we probe the molecular basis for cell plasticity in the preimplantation embryo in the context of regulative development. We find that cells of the early PrE are sufficient to regenerate both Epi and TE, forming intact blastocysts that upon implantation undergo normal postimplantation development until at least embryonic day 6.5 (E6.5). We recapitulate this plasticity *in vitro* using nEnd and demonstrate that OCT4/PDGFRA co expressing cells are uniquely competent to form Epi and TE upon targeted differentiation and chimera formation. Plasticity in early extra-embryonic endoderm is supported by JAK/STAT signalling and occurs via Oct4-mediated priming of transcriptionally quiescent enhancers.

## Results

### The E3.5 PrE maintains multilineage plasticity

The regulative properties of the mammalian preimplantation embryo have been shown by numerous grafting experiments, including the demonstration that the TE can be regenerated from an isolated ICM at the mid-blastocyst stage^4^. Given that lineage specification in the mouse ICM proceeds progressively between E3.0-3.5 without lineage fluctuations^13^, we asked which lineage is responsible for this plasticity. We converted putative ICMs of 8-cell embryos into either PrE or Epi by treatment with FGF4 and the MEK inhibitor PD0329501 (PD03), respectively, for 24h^29, 30^, after which we removed the TE by immunosurgery (condition D and F)^31^ (Fig. 1A). To confirm that treated ICMs were homogenously PrE or Epi, we quantified the proportion of cells expressing the PrE-marker GATA6 and the Epi-marker NANOG. In FGF4-treated embryos (condition D), we found that 76.9% were entirely single positive for GATA6, where the remaining 22.1% of embryos contained a few cells expressing high levels of GATA6 with low levels of NANOG (Fig. 1B; Suppl. 1A, B), consistent with previous studies^32^. There were no embryos containing NANOG single positive cells. In PD03-treated embryos (condition F), 81.2% of these were single positive for NANOG (Suppl. 1A, C, D). Quantification of CDX2 expression in the isolated ICMs across all conditions demonstrated no CDX2 positive cells following immunosurgery in the majority of embryos and the few that did (14%), contained ≤2 CDX2 positive cells with significant damage to their membrane integrity, indicating that these are likely not functional (Suppl. 1E). Following 48h of recovery, the TE was reconstructed *de novo* in 79.2% of control embryos (condition C) and 63.6% of embryos treated with FGF4 (condition E), but only 29% of embryos treated with PD03 (condition G) based on quantification of immunostaining for the TE marker CDX2 (Fig. 1B, C; Suppl. 1A).

**Figure 1:**
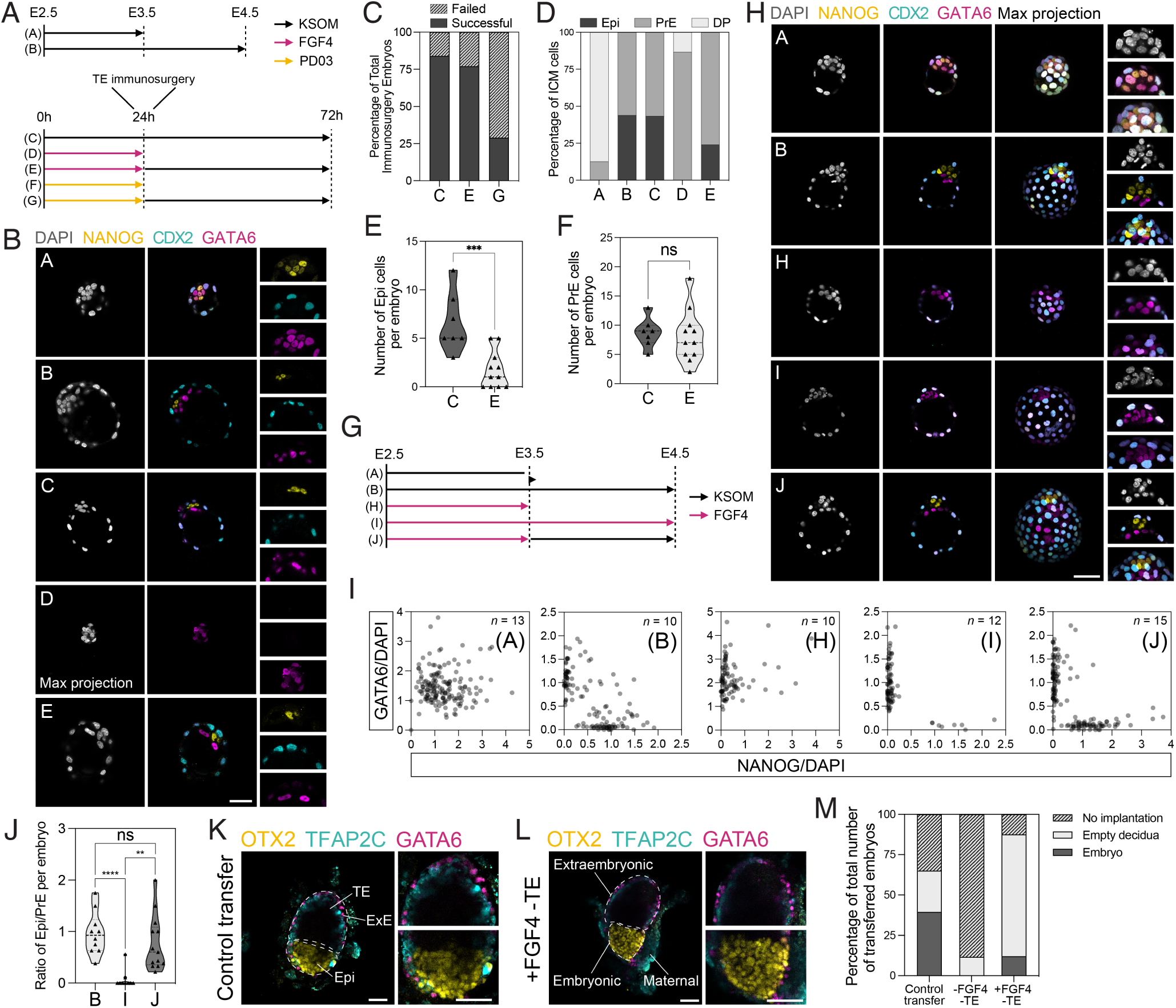
The E3.5 PrE reconstructs all embryonic and extra-embryonic lineages following perturbation. **(A)** Schematic of treatment regimen that 8-cell embryos were subjected to in **B**-**F**; Suppl. 1A-E. **(B)** Representative immunostaining of embryos for the indicated markers including DAPI for visualisation of nuclei, imaged by confocal microscopy. **(C)** Embryo survival following immunosurgery. **(D)** ICM lineage allocation in control and treated. **(E, F)** Allocation of **E** Epi and **F** PrE cells per ICM in control and treated embryos. **(G)** Schematic of treatment regimen that 8-cell embryos were subjected to in **H**-**J**; Suppl. 1F. **(H)** Representative immunostaining of E4.5 embryos for the indicated markers including DAPI for visualisation of nuclei, imaged by confocal microscopy. **(I)** Single cell quantification of embryos for NANOG and GATA6 immunostaining normalized to DAPI. *n* values indicate total number of embryos quantified. **(J)** Ratio of Epi to PrE cells per embryo. **(K, L)** Representative immunostaining of **K** control and **L** treated E6.5 embryos for indicated markers, where dashed white lines indicate embryonic and extra-embryonic regions. **(M)** Embryo survival and developmental progression across indicated treatments. *n* values for **B**-**F** in Suppl. 1A, **H-J** in Suppl. 1G and **K-M** in Suppl. 1I. Scale bars represent 50µm.

While all control embryos were able to form a blastocyst containing Epi, PrE and TE at normal proportions, marked by NANOG, GATA6 and CDX2 expression, this was the case for only 63.6% of the FGF4-treated embryos (condition E), where 36.4% of these formed blastocysts containing only CDX2 and GATA6 positive cells (Fig. 1B, D; Suppl. 1F). When quantifying the allocation of ICM cells in the embryos that did successfully specify an Epi, FGF4-treated embryos also contained significantly less Epi cells per ICM compared to control embryos, while establishing normal PrE cell numbers (Fig. 1E, F). This suggests that in a context where a PrE cell within the ICM is faced with the decision become either Epi or TE, it prioritizes TE.

To determine whether this deficiency in Epi regeneration is based on the competence of the PrE or the limited capacity of these cells to regenerate both Epi and TE lineages simultaneously, we again treated 8-cell embryos with FGF4 for 24h and asked whether these ICMs containing only PrE would efficiently generate Epi in the context of an intact TE when released from FGF4 stimulation (Fig. 1G). We found that the majority of embryos (89%) contained NANOG single positive cells after treatment, and the total levels of GATA6 and NANOG expression was similar to the control embryos (condition J) (Fig. 1H, I; Suppl. 1G). Indeed, when looking at the ratio of Epi/PrE cells per embryo, the FGF4-treated embryos were not significantly different to the controls (Fig. 1J). Transfer of these embryos to pseudopregnant mice produced live births (Suppl. 1H), overall confirming that the E3.5 PrE maintains competence for not only Epi differentiation, but complete development to term.

Having established that in a setting where an immunosurgery-isolated ICM is composed entirely of GATA6 positive cells, the PrE is able to specify both TE and Epi, we then reasoned that the PrE prioritizes making a functional TE first to ensure implantation success, after which specification of Epi will follow at a delayed rate. This is in agreement with the Epi requiring just four pluripotent cells to proceed with normal development^33^. To test the functionality of the *de novo* reconstructed TE, we transferred treated embryos that successfully reconstructed a blastocyst to pseudopregnant mice until E6.5, together with immunosurgery embryos without FGF4 treatment and normal E3.5 blastocysts (Suppl. 1I). While the reconstructed embryos without FGF4 treatment produced only a single empty decidua, the treated embryos underwent extensive decidualization based on morphology and immunostaining for the lineage-specific markers OTX2 (Epi), GATA6 (VE/PE) and TFAP2C (extra-embryonic ectoderm) (Fig. 1K-M; Suppl. 1J, K), demonstrating the remarkable plasticity of the early PrE to re-establish normal development.

### Naïve extra-embryonic endoderm creates a pluripotent niche upon self-organisation

To recapitulate the dynamic nature of the early PrE *in vitro*, we exploited a double reporter ESC line for both endoderm and Epi, GATA6-mCherry/SOX2-GFP (GCSG)^34^ and differentiated these to PrE (Suppl. 2A, B). After fluorescence activated cell sorting (FACS) for GATA6-mCherry single positive cells, these were expanded as nEnd in RPMI-based defined medium containing Activin A, CHIR99021 and LIF (RACL)^28, 35, 36^. Throughout nEnd culture, we sporadically observed spheroid aggregates arising from the underling the endodermal monolayer (Fig. 2A), that based on fluorescent imaging were composed of an outer layer of GATA6-mCherry positive cells and an inner core of SOX2-GFP/OCT4 positive cells (Fig. 2B). To confirm that these aggregates were spontaneously arising entirely from GATA6-mCherry-expressing nEnd, we isolated GATA6-mCherry single positive cells by FACS (Suppl. 2C) and found that a SOX2-GFP single positive population arose *de novo* within 48-72h after seeding (Fig. 2C-E). This was also observed when isolating nEnd from a wild type E14JU cell line based on PDGFRA-APC expression, from which a PECAM-FITC single positive population arose within the same window of time (Suppl. 2D). When replating the *de novo* SOX2-GFP cells, we found that these readily differentiated back towards PrE based on upregulation of GATA6-mCherry (Fig. 2E). We confirmed that these were not a product of contaminating ESCs left over from differentiation by performing consecutive rounds of FACS to isolate pure populations of GATA6-mCherry nEnd (Fig. 2F).

**Figure 2:**
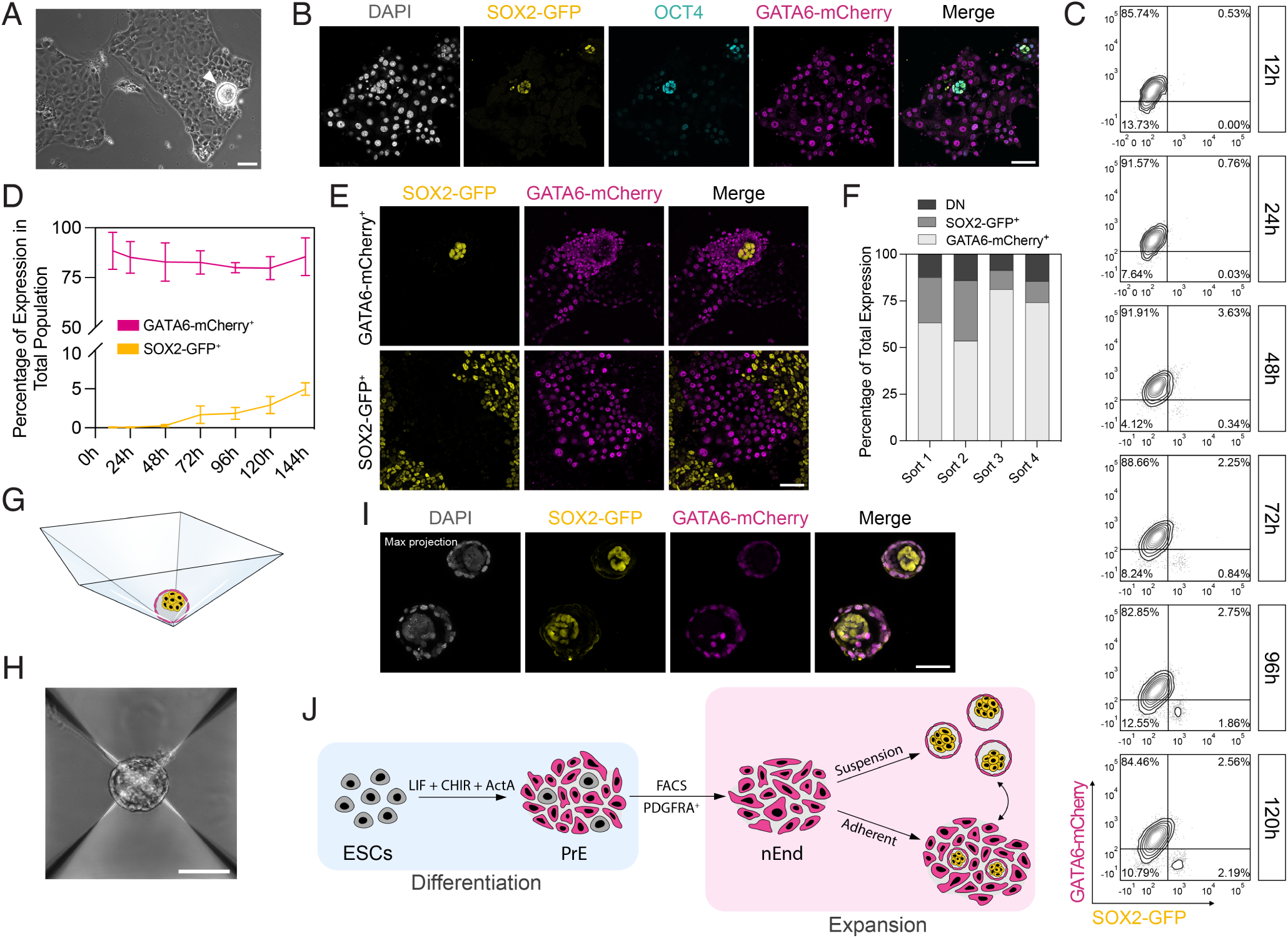
GATA6-expressing nEnd spontaneously undergoes de-differentiation to a SOX2-expressing Epi-like cell type. **(A)** Brightfield image of nEnd with aggregate emerging from monolayer (white arrowhead). **(B)** Immunostaining of GCSG nEnd for OCT4 including DAPI for visualisation of nuclei, imaged by confocal microscopy. **(C)** Representative flow cytometry contour plots of GCSG nEnd 12-120h following FACS for GATA6-mCherry^+^/SOX2-GFP^-^ cells (*n* = 4 biological replicates). Bottom left quadrant indicates gating based on a negative control. **(D)** Quantification of total expression by flow cytometry of GCSG nEnd 12-144h following FACS for GATA6-mCherry^+^/SOX2-GFP^-^ cells (*n* = 4 biological replicates). **(E)** Representative immunofluorescence imaging by confocal microscopy of GCSG nEnd 7 days following isolation by FACS for either GATA6-mCherry^+^ or SOX2-GFP^+^ cells. **(F)** Quantification of percentage total expression of GCSG nEnd subpopulations across 4 consecutives rounds of FACS for GATA6-mCherry^+^ nEnd. Each timepoint was collected 5 days after seeding. **(G)** Illustration of 3D nEnd cultured in an AggreWell with magenta endodermal outside cells and yellow pluripotent inside cells. **(H)** Brightfield image of 3D nEnd cultured in an AggreWell. **(I)** Representative immunofluorescence imaging by confocal microscopy of GCSG 3D nEnd cultured in AggreWell’s. **(I)** Immunofluorescence image of GCSG 3D nEnd imaged by confocal microscopy. **(J)** Schematic of nEnd culture system. ESCs are differentiated towards PrE, whereafter FACS for PDGFRA-APC expression, these cells are expanded as either nEnd in adherent culture or 3D nEnd in suspension culture. Scale bars represent 100µm in **A, H**; 50µm in **B**, **E**, **I**.

As these de-differentiation events were difficult to predict in adherent culture, we asked whether we could generate reproducible structures in suspension culture that would support the transition between GATA6-mCherry positive nEnd to a SOX2-GFP Epi-like population using AggreWell’s (Fig. 2G), referred to as 3D nEnd. In this context, we observed the formation of spheroid structures with GATA6-mCherry positive outside cells and SOX2 GFP positive inside cells abutted to a cavity (Fig. 2H, I; Suppl. 2E, F). To determine whether a similar nEnd culture could be captured directly from the mouse embryo, we isolated E3.5 blastocysts, removed the TE by immunosurgery and placed the isolated ICMs into RACL medium (Suppl. 2G). During initial passages, these cells were homogenously PDGFRA-APC positive (Suppl. 2H), however upon culture in AggreWell’s, embryo-derived nEnd readily formed 3D nEnd with a GATA6 positive outer layer and NANOG/OCT4 positive de differentiated core (Suppl. 2I). Taken together, nEnd culture is dynamically heterogenous, containing both committed and uncommitted extra-embryonic endoderm, with a subpopulation capable of self-organisation into delaminated Epi and PrE-like cell types (Fig. 2J).

### Heterogeneity in nEnd is marked by a population expressing Oct4 with enhanced developmental potential

During preimplantation development, OCT4 is initially expressed in all cells of the ICM, including the early PrE, where it functions as an essential mediator of endoderm specification^9, 18–20, 37^. OCT4 reaches its peak expression in the nascent PrE immediately following its segregation from the Epi and is mostly lost in the PrE by the late blastocyst stage^18^. To further map *Oct4* mRNA expression across PrE maturation from scRNA-seq of the preimplantation embryo^38^ (Suppl. 3A), referred to as Nowotschin 2019, we employed RNA velocity using scVelo (Fig. 3A)^39–41^. While Nanog is rapidly downregulated in the first PrE progenitors, *Oct4* follows *Gata6* expression throughout the early phase of PrE specification. *Oct4* is then gradually downregulated concurrently with upregulation of the late PrE marker *Gata4*.

**Figure 3:**
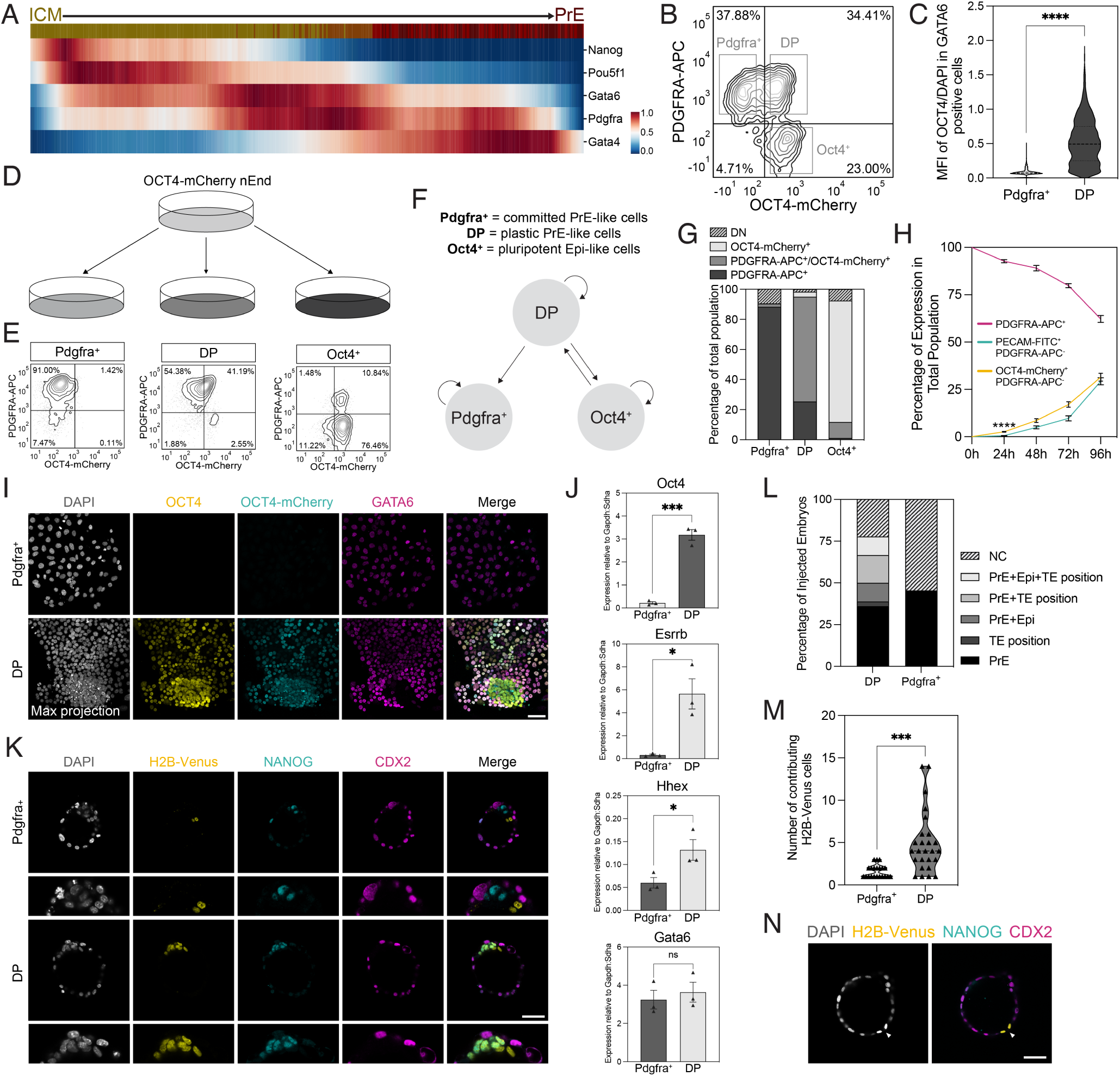
OCT4/PDGFRA co-expressing nEnd represents early uncommitted PrE capable of multilineage differentiation. **(A)** Heatmap of scaled expression of indicated genes from scRNA-seq of the mouse preimplantation embryo^38^ across scVelo-defined latent time. Top bar shows progression from E3.5 ICM (dark yellow) to E4.5 PrE (dark red). **(B)** Flow cytometry contour plot of OCT4-mCherry nEnd stained for PDGFRA-APC. Grey boxes highlight Pdgfra^+^, DP and Oct4^+^ populations, with bottom left quadrant indicating gating based on a negative control. **(C)** Mean fluorescence intensity (MFI) of OCT4 expression by immunostaining normalized to DAPI (*n* = 1495 (left) and 1467 (right) cells). **(D)** Schematic of OCT4-mCherry nEnd subpopulation re-plating experiment corresponding to **E**. **(E)** Representative flow cytometry contour plots of Pdgfra^+^, DP and Oct4^+^ nEnd 5 days following FACS for PDGFRA-APC^+^ cells, where bottom left quadrant indicates gating based on a negative control. **(F)** Schematic of dynamic equilibrium established in nEnd culture. **(G)** Quantification of **D**, **E** for subpopulation composition in total population for Pdgfra^+^, DP and Oct4^+^ nEnd (DN = double negative). **(H)** Quantification of total expression by flow cytometry of OCT4-Cherry nEnd 0-96h following FACS for PDGFRA-APC^+^ cells performed in Suppl. 3E. **(I)** Immunostaining of OCT4-mCherry nEnd for indicated markers including DAPI for visualisation of nuclei 5 days following FACS for PDGFRA-APC^+^ cells. **(J)** RT-qPCR of Pdgfra^+^ and DP nEnd isolated by FACS for indicated markers. **(K)** Representative immunostaining of E4.5 embryos injected with either OCT4-mCherry-H2B-Venus DP (top) or Pdgfra^+^ (bottom) nEnd at the 8-cell stage for indicated markers including DAPI for visualisation of nuclei, imaged by confocal microscopy. **(L)** Allocation of lineage contribution of OCT4-mCherry-H2B-Venus DP (*n* = 36) and Pdgfra^+^ (*n* = 44) nEnd in chimera embryos (NC = no contribution). **(M)** Total number of OCT4-mCherry-H2B-Venus DP and Pdgfra^+^ cells contributing to each embryo (*n* values as in **L**). **(N)** Representative immunostaining of OCT4-mCherry-H2B-Venus DP nEnd chimera embryo for indicated markers including DAPI for visualisation of nuclei, imaged by confocal microscopy. Scale bars represent 50µm.

Consistent with nEnd trapping a preimplantation PrE-like state, we found that OCT4 was heterogeneously expressed throughout GATA6 positive nEnd culture organised into distinct patches (Suppl. 3B). To explore nEnd dynamics, we employed an OCT4-mCherry protein-based reporter cell line^42^ and by flow cytometry, observed three distinct populations based on co-expression with PDGFRA-APC: PDGFRA-APC single positive (Pdgfra^+^), PDGFRA-APC/OCT4-mCherry double positive (DP) and OCT4-mCherry single positive (Oct4^+^) (Fig. 3B, C; Suppl. 3C). Here, Pdgfra^+^ and DP comprise the endodermal portion of nEnd culture, while Oct4^+^ represent the de-differentiated pluripotent cells based on co expression with PECAM-FITC (Suppl. 3D). To determine the dynamic properties of the three populations, we isolated these by FACS and re-plated them individually in adherent nEnd culture (Fig. 3D). After 120h, DP nEnd were able to give rise to themselves, as well as the other two populations, whereas Pdgfra^+^ cells remained PDGFRA-APC single positive, with a small population becoming double negative (DN) (Fig. 3E-G). Oct4^+^ nEnd were similarly able to self-renew, in addition to undergo differentiation towards DP cells (Fig. 3E-G). When re plating DP cells in adherent nEnd culture and monitoring expression at 24h intervals for 96h, an Oct4^+^ population precedes PECAM-FITC expression, which arises after 24h and 48h, respectively (Fig. 3H; Suppl. 3E). DP and Pdgfra^+^ nEnd are also morphologically distinct and divide at different rates. Although they share similar nuclear/cytoplasmic ratios (0.63 and 0.67 for DP and Pdgfra^+^, respectively), DP cells form compact colonies with overall cell and nuclei size that are significantly smaller compared to Pdgfra^+^ cells, which instead form large epithelial sheets (Suppl. 3F-H) and have a doubling time that is 1.3x faster than Pdgfra^+^ nEnd (Suppl. 3I).

In adherent nEnd culture, DP nEnd forms OCT4 single positive dome-shaped colonies surrounded by endodermal monolayers containing a mixture of OCT4/GATA6 double positive cells and GATA6 single positive cells, whereas Pdgfra^+^ cells form only a GATA6 single positive monolayer (Fig. 3I). Similarly, upon seeding in 3D nEnd culture, only DP cells formed spheroids with both a GATA6 positive outer layer and an OCT4/NANOG double positive inner core, whereas Pdgfra^+^ cells formed GATA6 single positive hollow spheres (Suppl. 3J). RT-qPCR revealed that DP cells express significantly higher levels of *Oct4*, *Esrrb* and *Hhex* compared to Pdgfra^+^ cells, pointing towards an earlier uncommitted PrE identity, while expressing similar levels of *Gata6* (Fig. 3J).

To benchmark nEnd culture within the developmental progression of extra-embryonic endoderm, we generated XEN cells from 2iLIF ESCs^26, 43^ (Suppl. 4A-C). We observed that XEN cells appeared refractile by morphology (Suppl. 4C) and expressed GATA6, but not OCT4^27^ (Suppl. 4D) and did not readily undergo de-differentiation, based on PECAM-FITC expression by flow cytometry (Suppl. 4E). When transferring XEN cells to RACL in both adherent and suspension culture, XEN cells failed to upregulate markers unique to nEnd culture (Suppl. 4E-G), including *Oct4* and *Esrrb*, instead expressing high levels of the VE marker *Afp* (Suppl. 4G). Oct4 is not expressed in XEN cells^26, 27^, and we failed to observe induction of Oct4 upon transfer to nEnd medium, further suggesting that XEN cells represent a committed extra-embryonic endodermal cell type.

### nEnd exhibits enhanced lineage competence

To test the competence of different nEnd subpopulations, we reintroduced these into normal development in chimera experiments. To visualize the fate of the injected cells, we labelled the OCT4-mCherry reporter with a constitutive lineage marker, termed OCT4 mCherry-H2B-Venus. 8-cell host embryos were injected with 3 DP or Pdgfra^+^ cells and allowed to develop for 48h until E4.5. DP cells efficiently contributed to 77.8% of injected embryos, with a high degree of proliferation, compared to just 45.5% of embryos injected with Pdgfra^+^ cells (Fig. 3K-M). DP cells were found populating all three lineages, while Pdgfra^+^ cells were only found in the PrE (Fig. 3K, L). While DP cells localised and appeared morphologically indistinguishable to TE, they did not yet upregulate CDX2, a transcription factor important for the TE transcriptional programme, and were therefore termed TE position cells^44^ (Fig. 3L, N).

While our observations regarding the potential of nEnd to initiate TE specification were inconclusive, a recent study found TSCs could be derived from an extra-embryonic endoderm-like intermediate^45^. We therefore asked whether nEnd could differentiate into TSCs *in vitro*. PDGFRA-APC single positive cells were isolated by FACS and cultured in MEF-conditioned medium containing FGF4 and heparin for 7 days. These produced a heterogenous population by morphology (Suppl. 5A), containing cells that were both CDX2 positive/BRACHYURY negative, suggesting a TE-like cell type, and CDX2/BRACHYURY double positive, suggesting a mesodermal-like cell type (Suppl. 5B). To distinguish TE from mesodermal cell types in culture, we performed scRNA-seq on nEnd sorted for PDGFRA APC by FACS and cultured in either RACL for 96h or TSC medium for 168h (Suppl. 5C). After quality control, the dataset contained 11,194 cells from both nEnd and TSC conditions, detecting 32,285 genes that upon sub-clustering and dimensionality reduction produced 12 clusters visualised by Uniform Manifold Approximation Projection (UMAP)^46^ (Suppl. 5D, E). These were annotated based on marker expression, with cells cultured in RACL as nEnd 1-5 and those cultured in TSC conditions as XEN-like 1-5, TE-like and mesoderm-like (Mes-like) (Suppl. 5F, G). nEnd 1-4 expressed high levels of Gata6 and Pdgfra, demonstrating endodermal identity, and nEnd 5 appeared to be the de-differentiated Epi-like cells based on expression of various pluripotency genes and the absence of PrE markers. XEN-like clusters 1-5 expressed high levels of pan-endodermal and PE markers, such as Gata6, Plat and Sparc, with few cells expressing the VE marker Afp. The appearance of a XEN-like phenotype in TSC medium is not surprising, as there are many similarities between TSC and XEN medium (see Methods and Materials). While both the TE-like and Mes-like clusters expressed Cdx2 and Gata3, only the Mes-like cells expressed the mesoderm-specific markers T, Mixl1 and Mesp1, with TE-like cells expressing significantly higher levels of the TE-specific markers Hand1, Krt8 and Krt18 (Suppl. 5H-J). When comparing differentially expressed genes (DEGs) between TE-like and Mes-like clusters to the Nowotschin 2019 dataset^38^, we found that the TE-like populations exhibited enriched gene expression reminiscent of the E3.5 and E4.5 TE, while the Mes-like population did not share gene enrichment with any of the *in vivo* cell type in this dataset (Suppl. 5K). This analysis demonstrates that nEnd is capable of differentiating towards TE-like cells that resemble the preimplantation TE *in vivo*.

### OCT4-expressing endoderm undergoes commitment in the absence of JAK/STAT signalling

We previously reported that LIF specifically supports PrE during preimplantation development by priming cells towards a PrE fate through the JAK/STAT pathway^47^, in addition to its role in supporting ESC self-renewal^48, 49^ and ICM identity *in vivo*^50^. As LIF is a key component of nEnd culture medium, we assessed the influence of LIF withdrawal or culture in a JAK inhibitor (JAKi)^47, 51^. We observed that prolonged culture in the absence of LIF or inhibition of JAK/STAT signalling resulted in a loss of the DP population through a significant reduction in phosphorylation of STAT3 and OCT4-mCherry expression (Fig. 4A-C; Suppl. 6A, B). In a complete block to JAK/STAT activity, de-differentiation at level of NANOG expression is lost (Fig. 4A). Similarly, flow cytometry analysis revealed a significant increase in Pdgfra^+^ cells at the expense of the DP population (Fig. 4D, E; Suppl. 6C, D).

**Figure 4:**
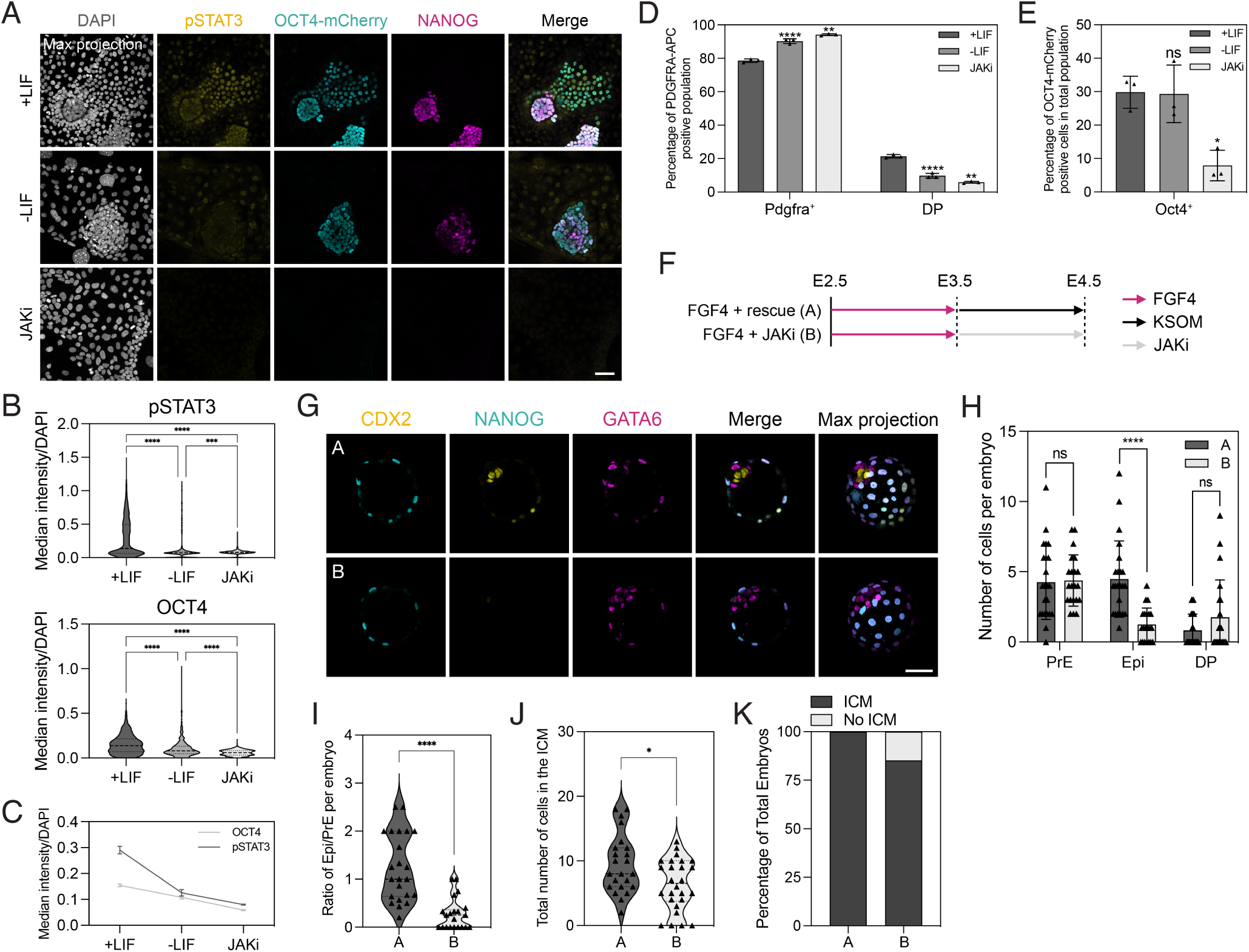
JAK/STAT signalling is required to maintain an early uncommitted PrE cell type *in vitro* and *in vivo*. **(A)** Representative immunostaining of OCT4-mCherry nEnd in control and treated conditions for indicated markers including DAPI for visualisation of nuclei, imaged by confocal microscopy. Cells cultured in conditions for 2 passages and single cell quantification performed in Suppl. 6B. **(B)** Quantification of median fluorescence intensity normalized to DAPI for **A**, where +LIF: *n* = 1629 cells; -LIF: *n* = 873 cells; JAKi: *n* = 726 cells based on Suppl. 6B. **(C)** Total median fluorescence intensity normalized to DAPI for **A**. **(D, E)** Total expression by flow cytometry of **D** PDGFRA-APC and **E** OCT4-mCherry for control and treated conditions in **A**. **(F)** Schematic of treatment regimen that 8-cell embryos were subjected to in **G**-**K**. **(G)** Representative immunostaining of E4.5 embryos for the indicated markers including DAPI for visualisation of nuclei, imaged by confocal microscopy. Expression of GATA6 and NANOG was used for analysis performed in **H**-**K**. **(H)** ICM lineage allocation in treated embryos. **(I)** Ratio of Epi to PrE cells per embryo in treated embryos. **(J)** Total number of cells within the ICM of treated embryos. **(K)** Quantification of presence or absence of an ICM in treated embryos. *n* values for treatment regimen outlined in **A** shown in **G**-**K** are FGF4+rescue (A): *n* = 23; FGF4+JAKi (B): *n* = 26 embryos. Scale bars represent 50µm.

To discern the role JAK/STAT signalling PrE plasticity *in vivo*, we first cultured 8-cell embryos in FGF4 for 24h to convert the ICM to PrE, after which the embryos were transferred either to control media (FGF4+rescue) or media containing 1µM JAKi (FGF4+JAKi) for an additional 24h (Fig. 4F). Rescue embryos were able to correctly allocate both the Epi and PrE lineage, while we observed that the JAKi-treated embryos either contained mostly only PrE or no ICM at all (Fig. 4G-K), suggesting that sustained JAK/STAT signalling is required to forestall differentiation of PrE, such that it retains competence for conversion towards Epi *in vivo*, as well as *in vitro*.

### nEnd contains a unique subpopulation enriched for OCT4 targets

To resolve nEnd heterogeneity and construct a transcriptional map of the subpopulations observed in nEnd culture, we performed scRNA-seq on nEnd in adherent culture, 3D nEnd in AggreWell’s and 2iLIF ESCs (Suppl. 7A). Quality control produced a dataset comprising of 14,788 cells and 32,285 genes from which 14 clusters were visualised by UMAP (Fig. 5A; Suppl. 7B). 2iLIF ESCs produced 5 clusters as well as the majority of a small cluster (137 cells) characterized by 2C gene expression^52^, which was annotated as 2C (Suppl. 7C). Expression of *Pdgfra* and *Gata6* was restricted to nEnd and 3D nEnd clusters, except the de-differentiated cluster expressing pluripotency markers, including *Nanog* and *Oct4*, annotated as Oct4^+^ (cluster 6) (Fig. 5B). Within the *Pdgfra*/*Gata6* expressing clusters, we identified cluster 7 and 14 as containing cells also expressing *Oct4*. As cluster 14 expressed high levels of apoptotic markers (Suppl. 7D), with enrichment for gene ontology terms related to cell death (Suppl. 7E), we excluded it from further analysis and refer to it as Apoptotic. Cluster 7 was therefore annotated as DP and the remaining nEnd and 3D nEnd clusters were annotated as Pdgfra^+^ 1-5 (Fig. 5A). These annotations agree with an nEnd culture that largely contains endoderm with a subpopulation of cells that have undergone de differentiation to an Epi-like identity. Principal component analysis (PCA) showed that the first principal component (PC1) separated all 2iLIF ESCs and a subset of nEnd and 3D nEnd cells from the bulk endoderm populations, with a branch of cells connecting the two (Suppl. 7F). This branch consists of the Oct4^+^ cluster bridging the pluripotent 2iLIF clusters to the endodermal Pdgfra^+^ clusters, with the DP cluster being an anchoring point for the latter.

**Figure 5:**
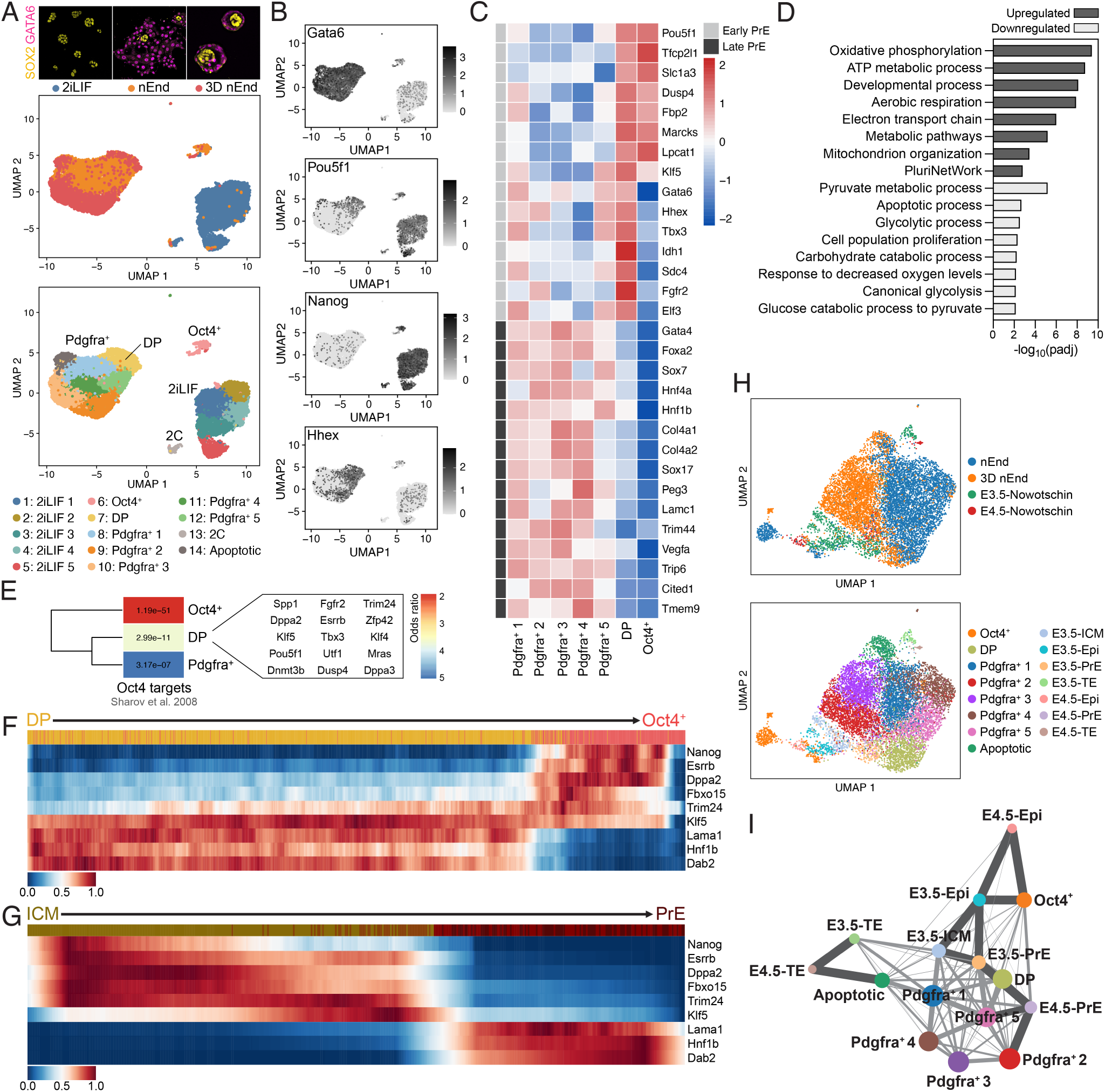
scRNA-seq of nEnd contains subpopulations that mirror stages of mouse preimplantation development. **(A)** UMAP dimensional embedding of 14,788 2iLIF, nEnd and 3D nEnd cells by scRNA-seq. Top: colouring based on culture condition shown above including representative immunostaining for SOX2 and GATA6; bottom: colouring based on Louvain clustering. **(B)** UMAP dimensional embedding showing single cell expression of indicated markers. **(C)** Heatmap of candidate lineage markers in log2 normalized clustered data for early and late PrE *in vivo*^38^ defined in Suppl. 7H. Scaled by row. **(D)** Gene ontology analysis of selected biological processes of up and downregulated genes in the DP nEnd cluster. **(E)** Comparative gene expression overlap analysis of Oct4^+^, DP and Pdgfra^+^ 1-5 clusters with 700 downstream targets of Oct4^59^. Colour scale is based on odds ratio and grids labelled with *p*-value indicating significance of the respective odds ratio. **(F, G)** Heatmap of scaled expression of indicated genes from scRNA-seq of **F** DP and Oct4+ nEnd clusters *in vitro* and **G** ICM to PrE *in vivo*^38^ across scVelo-defined latent time. Top bar shows progression from **F** DP (yellow) to Oct4^+^ (pink) and **G** E3.5 ICM (dark yellow) to E4.5 (dark red). Genes are listed in the same order for both **F** and **G**. **(H)** UMAP dimensional embedding of integrated *in vitro* nEnd and 3D nEnd (this study) and *in vivo*^38^ datasets using SCVI^56^. Top: colouring based on culture condition (nEnd, 3D nEnd) or developmental stage (E3.5, E4.5). Bottom: colouring based on Louvain clustering. **(I)** PAGA of integrated scRNA-seq dataset shown in **H**. Dark grey lines indicates highly connected regions and light grey lines indicate regions with lower confidence.

When probing the expression of a panel of lineage-specific markers for Epi, ICM, PrE, VE and PE across all nEnd/3D nEnd clusters (Pdgfra^+^ 1-5, DP and Oct4^+^) (Suppl. 7G), we found Epi-related markers specifically upregulated in the Oct4^+^ cluster and PrE, VE and PE marker expression at varying degrees in Pdgfra^+^ clusters 1-5. The DP cluster has a unique profile, expressing canonical PrE genes while maintaining modest levels of select ICM and pluripotency genes. When expanding the panel of PrE markers to those specific to early and late PrE development *in vivo*^38^ (Fig. 5C; Suppl. 7H), the nEnd clusters partitions into two, where DP falls into early PrE and the remaining nEnd clusters resemble late PrE. As XEN cells have been identified as an intermediate in the reprogramming of somatic cells to pluripotency^53, 54^, we considered whether nEnd and XEN intermediates represent a common cell type. We exploited scRNA-seq of chemical somatic cell reprogramming towards iPSCs^55^ and integrated our datasets using SCVI^56^. We quantified cell-type connectivity with partition based graph abstraction (PAGA)^57^ (Suppl. 7I) and found that the biproduct XEN-like population derived from TSC differentiation of nEnd (Suppl. 5) strongly aligns with the XEN like intermediates identified in reprogramming, while the nEnd subpopulations closer resemble the cell types arising later in the trajectory towards pluripotency.

We then performed gene ontology analysis of the top 300 DEGs based on comparison of DP vs Pdgfra^+^ 1-5 clusters (Suppl. 8A). We found enrichment of metabolic terms associated with oxidative phosphorylation, mitochondrial activity, and pluripotency in DP cells, while Pdgfra^+^ cells were associated with glycolytic processes (Fig. 5D). To verify these observations *in vitro*, we cultured OCT4-mCherry nEnd sorted for PDGFRA-APC single positive expression and cultured these in either an inhibitor of glucose metabolism (2DG) or an inhibitor of CPT1A and fatty acid oxidation (Etomoxir). Here, we observed that increasing oxidative phosphorylation with 2DG resulted in a significant increase in DP cells, while an increase in glycolytic metabolism with Etomoxir led to an increase in Pdgfra^+^ cells (Suppl. 8B, C).

We found many of the genes that were differentially upregulated in the DP cluster are known interactors and downstream targets of OCT4^58, 59^. To establish the significance of this, we compared all differentially upregulated genes in Oct4^+^, DP and Pdgfra^+^ 1-5 clusters to 700 functional downstream target genes of OCT4 identified by ChIP and microarray^59^ (Fig. 5E). The DP cluster positioned itself as an intermediate between Pdgfra^+^ 1-5 and Oct4^+^, where Oct4 targets were modestly enriched in the DP cluster and strongly enriched in the Oct4^+^ cluster. Of the 361 Oct4 targets enriched in Oct4^+^, 70 of these were common between Oct4^+^ and DP, such as *Oct4*, *Esrrb* and *Dppa2* (Suppl. 8D).

Having established that the Oct4^+^ de-differentiated cells are derived entirely from DP nEnd (Fig. 3D-F), we then asked what transcriptional changes are occurring between the DP and Oct4^+^ cluster. Based on DEGs, pluripotency and Epi markers are significantly upregulated in Oct4^+^ cells, while endoderm and extra-cellular matrix markers are significantly downregulated (Suppl. 8E), consistent with an endodermal cell type progressing towards a pluripotent Epi-like state.

To infer directionality to the transition of a PrE-to-Epi identity in our dataset, we employed RNA velocity^39, 41^ to visualise latent time by UMAP (Suppl. 8F, G). From this, we observed the single cell trajectory of gene expression from the DP to Oct4^+^ clusters and compared this to the transition from E3.5 ICM to E4.5 PrE^38^ for select downstream targets of Oct4^59^ (Fig. 5F, G). Here, we found the gene expression pattern to be directly inversely related to each other with early PrE markers, such as *Dab2* and *Klf5*, downregulated during the transition from DP to Oct4^+^ concurrent with upregulation of ICM/Epi markers, such as *Dppa2* and *Esrrb*. By contrast, these factors were reciprocally regulated in the ICM to PrE trajectory.

As DP nEnd exhibits a number of characteristics of early PrE, we directly compared this state to gene expression during PrE specification *in vivo*. We integrated our scRNA-seq dataset, excluding 2iLIF cells, with mouse preimplantation embryo stages E3.5 and E4.5^38^ using SCVI^56^ (Fig. 5H), and used PAGA^57^ to determine trajectory inference across all conditions. We found our DP cluster positioned directly between E3.5 and E4.5 PrE, while our Oct4^+^ cluster positioned between E3.5 and E4.5 Epi (Fig. 5I). Overall, this suggests DP nEnd is a good model for blastocyst stage PrE just prior to commitment^32, 60^.

### ESRRB is required downstream of OCT4 to support plasticity

To establish whether Oct4 is directly regulating plasticity in nEnd or merely a marker of early PrE, we performed degradation of Oct4 by siRNA and assessed whether this impairs plasticity (Fig. 6A). We observed a significant knockdown of *Oct4* mRNA (∼60%) in nEnd by RT-qPCR, as well as reduced expression of the Oct4 targets *Esrrb* and *Zfp42*, that are normally upregulated in de-differentiation (Fig. 6B). We similarly observed a significant increase in Pdgfra^+^ cells at the expense of the DP population by flow cytometry (Fig. 6C). ESRRB interacts with OCT4, where they are found in the same complex, as well as cooperatively regulating parts of the pluripotency network^58, 61^, and we recently found that Esrrb supports plasticity by safeguarding the decision point between Epi and PrE-like cells^40^. We probed the relationship between *Oct4* and *Esrrb* during de-differentiation from DP to an Oct4^+^ state by employing tetracycline-inducible Esrrb knockout (Esrrb^-/-^:tetON-Esrrb (EKOiE)) ESCs^62^. These were differentiated to PrE by inducing Esrrb expression for the first 48h, after which doxycycline (Dox) was removed for the duration of differentiation and subsequent nEnd expansion. EKOiE nEnd was sorted based on PDGFRA-APC expression by flow cytometry and seeded in the presence (+Dox) or absence (-Dox) of Dox for 7 days (Fig. 6D). We monitored the pluripotency marker PECAM and found that only +Dox nEnd was able to undergo de-differentiation (Fig. 6E). This was confirmed by immunofluorescence where only +Dox nEnd formed OCT4 single positive aggregates (Fig. 6F). While -Dox nEnd formed large epithelial sheets lacking ESRRB expression and points of de-differentiation, we did observe OCT4 co-expression with GATA6 throughout the culture (Fig. 6F, G). As OCT4 protein is significantly increased in GATA6^+^ nEnd +Dox compared to -Dox (Fig. 6H), we asked whether ESRRB is stabilising OCT4 protein or actively regulating its transcription. RT qPCR of PDGFRA-APC positive nEnd +Dox and -Dox revealed levels of *Oct4* and nascent *Oct4* were unchanged despite increased levels of *Esrrb* (Fig. 6I). This suggests that ESRRB acts downstream to stabilize OCT4, enabling the higher levels of protein expression required for de-differentiation and activation of the Epi programme.

**Figure 6:**
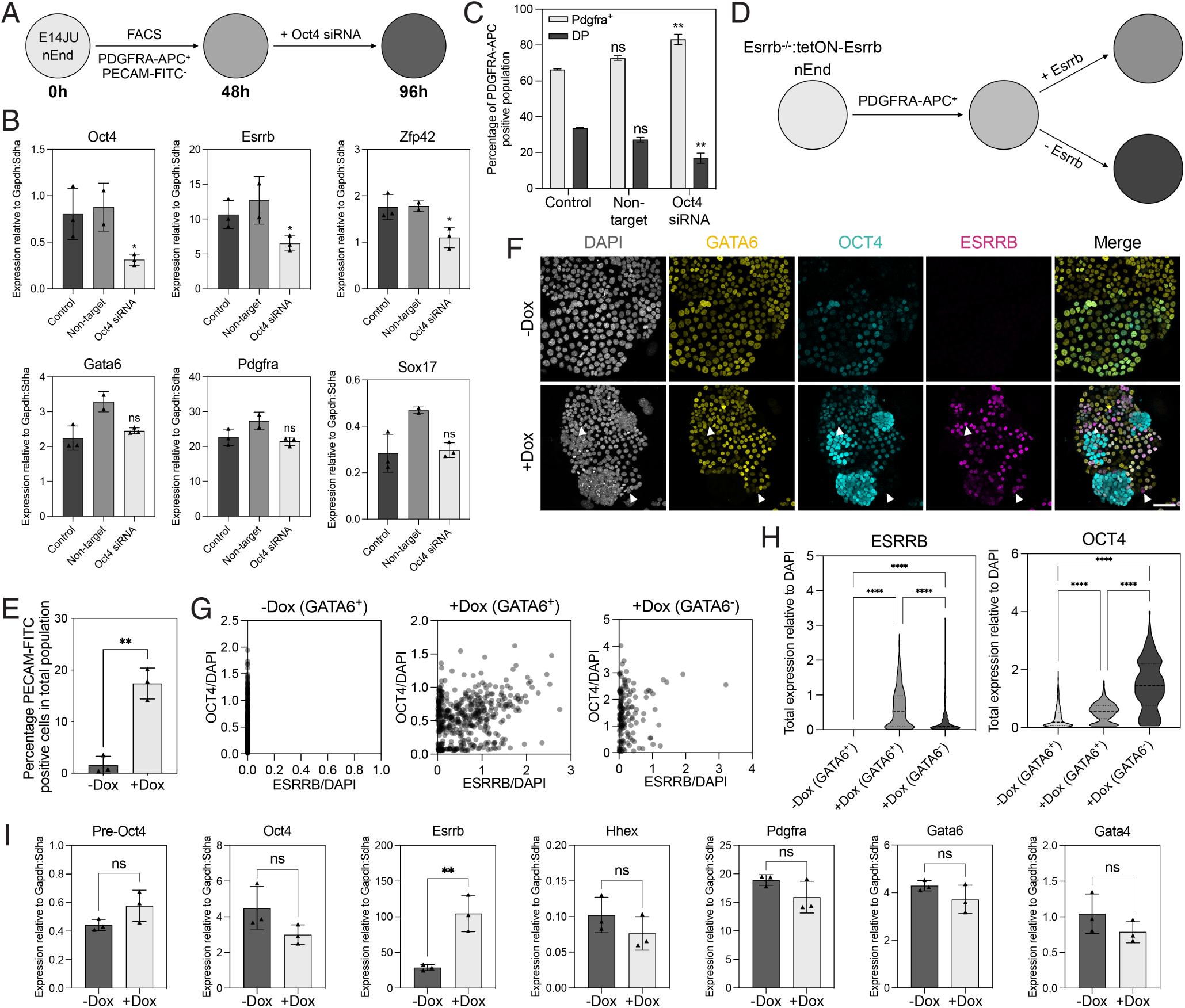
OCT4 is stabilised by ESRRB to enable de-differentiation of DP nEnd to an Epi-like cell type. **(A)** Schematic of Oct4 siRNA knockdown. E14JU nEnd were isolated by FACS for PDGFRA-APC expression, where after 48h in culture, the cells were treated with Oct4 siRNA or a non-targeting control siRNA for an additional 48h. **(B)** RT-qPCR of siRNA-treated nEnd for the indicated markers. **(C)** Quantification of proportions of DP and Pdgfra^+^ cell based on total PDGFRA-APC expression by flow cytometry for siRNA-treated nEnd. Statistics based on control vs non-target control and non-target control vs Oct4 siRNA. **(D)** Schematic of Esrrb induction with EKOiE nEnd. PDGFRA-APC^+^ EKOiE nEnd was isolated by FACS and cultured in +Dox (+Esrrb) or -Dox (-Esrrb) for 7 days. **(E)** Quantification of total expression of PECAM-FITC in EKOiE nEnd +/-Dox. **(F)** Representative immunostaining of EKOiE nEnd +/-Dox for indicated markers including DAPI for visualisation of nuclei, imaged by confocal microscopy. **(G, H)** Quantification of **F** for OCT4 and ESRRB in cells grouped by GATA6 expression for **G** single cell and **H** total expression. **(I)** RT-qPCR of PDGFRA-APC^+^ EKOiE cells +/-Dox isolated by FACS for indicated markers. Scale bar represents 50µm.

### The unique enhancer landscape in DP nEnd supports plasticity

To understand how Oct4-mediated plasticity in nEnd is supported at a transcriptional level, we performed cleavage under targets and tagmentation (CUT&Tag)^63^ to profile regulatory regions in Pdgfra^+^, DP and Oct4^+^ cells, focusing on key regulatory histone modifications, including H3K27ac and H3K4me1 for active and primed histone marks, respectively^64^, H3K27me3 for polycomb repressive complex 2 activity and enhancer priming^65^, and H3K9me3 for repression^66^. To annotate consensus peaks within our dataset, we incorporated enhancer peak sets for those specific to either pluripotency or nEnd^34^. Here, we observed that at pluripotency genes such as *Nanog*, Oct4^+^ cells were marked by both H3K27ac and H3K4me1, and DP cells by H3K4me1 only, with Pdgfra^+^ cells marked by neither (Suppl. 9A). By contrast endodermal loci, such as *Col4a1* and *Col4a2*, were found to be enriched in both H3K27ac and H3K4me1 marks at nEnd enhancers in DP and Pdgfra^+^ nEnd, but not in de-differentiated Oct4^+^ cells (Suppl. 9B). A global assessment of pluripotency and nEnd enhancers across all three populations for H3K4me1 revealed an enrichment of pluripotency enhancers in Oct4^+^ and an enrichment of nEnd enhancers in Pdgfra^+^, with DP positioning itself as an intermediate in both (Fig. 7A, B). While the deposition of H3K4me1 characterises priming of enhancer elements, bivalent regulatory regions with co-binding of histone marks such as H3K4me1 and H3K27me3 are associated with both active and repressive transcriptional outcomes^67^. To address how the enhancer state in DP nEnd cells could account for plasticity, we categorised our data based on regions containing H3K4me1 together with H3K27ac (active), H3K4me1 alone (primed), H3K4me1 together with H3K27me3 (bivalent) and H3K9me3 for repressed (Fig. 7C). When comparing pluripotent enhancers across these categories in Pdgfra^+^, DP and Oct4^+^ cells, we found that they were active in Oct4^+^ cells, maintained in either primed or bivalent states in the DP cells and repressed in committed Pdgfra^+^ nEnd (Fig. 7D). This contrasts nEnd enhancers, which were active and primed in Pdgfra^+^ nEnd, but bivalent in DP nEnd and inactive in Oct4^+^ cells (Fig. 7E).

**Figure 7:**
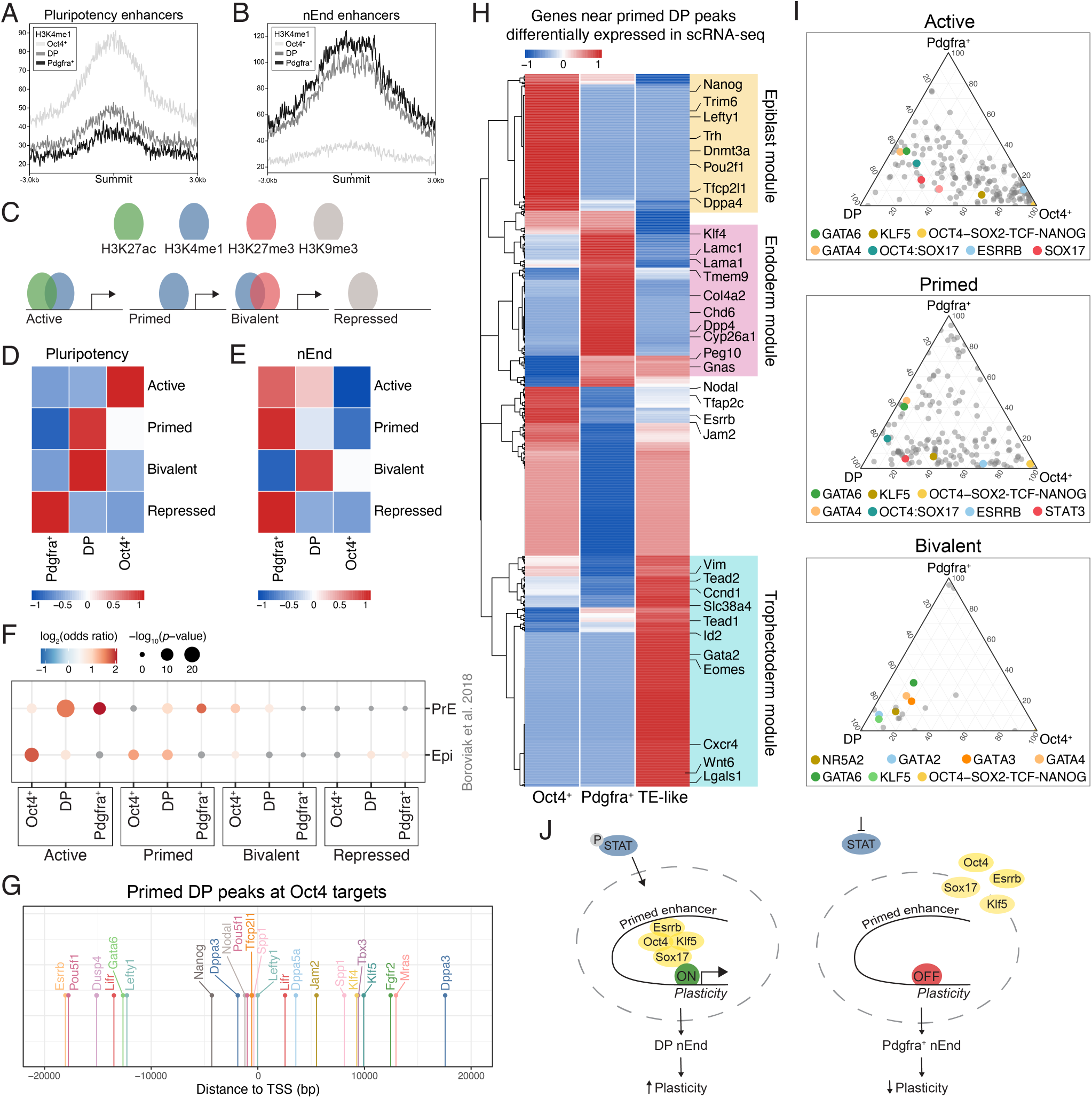
Status of lineage-specific regulatory elements supports DP nEnd plasticity and predicts future differentiation competence. **(A, B)** H3K4me1 occupancy at defined **A** pluripotency and **B** nEnd enhancers^34^ in Oct4^+^, DP and Pdgfra^+^ nEnd. **(C)** Schematic of histone mark categories for enhancer states. Active: H3K27ac and H3K4me1; Primed: H3K4me1 only; Bivalent: H3K4me1 and H3K27me3; Repressed: H3K9me3. **(D, E)** Heatmap of log_2_ normalized peaks overlapping with annotated **D** pluripotency and **E** nEnd enhancers. Scaled by row. **(F)** Comparative gene overlap analysis of CUT&Tag peaks annotated to the nearest TSS within promoter regions against DEGs from scRNA-seq data of the mouse preimplantation Epi and PrE *in vivo*^68^. Grey points represent *p* > 0.05. **(G)** Position of DP primed peaks annotated to the nearest TSS (±20kb) relative to selected downstream target genes of Oct4^59^ and Gata6. **(H)** Heatmap of scRNA seq data from log_2_ normalized DEGs in Oct4^+^, Pdgfra^+^ and TE-like clusters compared to DP nEnd subset for genes within ±20kb of primed DP peaks. Annotations include lineage specific genes for Epi, endodermal and TE lineages. **(I)** Ternary plots of motif enrichment across Oct4^+^, DP and Pdgfra^+^ nEnd in the active, primed, and bivalent categories. *p* < 0.05 for at least one population in every motif shown, where the axis represents -log10(*p*-value). Grey dots indicate all motifs and coloured dots highlight motifs of interest. **(J)** Proposed model for JAK/STAT and Oct4-mediated plasticity in enhancer priming.

To further understand the identity of nEnd subpopulations, we annotated all peaks to their nearest transcription start site (TSS) and compared these genes to DEGs from published scRNA-seq data of mouse preimplantation Epi and PrE *in vivo*^68^ (Fig. 7F). Genes regulated by Oct4^+^ and Pdgfra^+^ active and primed enhancers overlapped significantly with Epi and PrE, respectively. Genes regulated by active DP enhancers were enriched for PrE specific genes, supporting their canonical endoderm identity, while primed and bivalent DP enhancers were found to regulate both Epi and PrE genes. A conservative estimate of the genes regulated by primed DP enhancers based on a proximity of ±20kb, produces a list of numerous key OCT4 targets^59^, including *Dppa3*, *Lifr*, *Tfcp2l1*, *Esrrb* and *Oct4* itself (Fig. 7G), where priming at these genes correlate with their activation in de-differentiated active Oct4^+^ cells (Suppl. 9C). While Nanog is primed in DP nEnd only at its -45kb super enhancer (SE), both the -45kb and -5kb SE’s are activated in Oct4^+^ cells, where the -45kb SE also regulates nearby *Dppa3*^69, 70^ (Suppl. 9A). We also found that primed regulatory regions in DP nEnd were in proximity to a number of key TE determinants, such as *Tfap2c*, *Eomes*, *Gata2* and *Tead2*, but not the OCT4-repressed TE target *Cdx2* (Suppl. 9D). The availability of these enhancers is consistent with the ability of nEnd to differentiate towards a TE-like cell type (Suppl. 5). Moreover, we could use regulatory elements in primed DP to predict lineage trajectory through activation of these lineage-specific genes in transcriptomes for Pdgfra^+^ and Oct4^+^ clusters, as well as TE-like cells derived from TSC differentiation of nEnd (Fig. 7H; Suppl. 9E).

Gene ontology analysis of primed DP peaks revealed an enrichment in ICM, TE differentiation and blastocyst development (Suppl. 9F), supporting the notion that these enhancers are maintaining competence in this population to generate all three blastocyst cell types, if necessary. To explore the factors responsible for this priming event, we performed motif analysis on our categorised enhancer sets. The OCT4:SOX17 motif was present in active and primed DP nEnd, but not the OCT4-SOX2-TCF-NANOG motif, which was only present in Oct4^+^ cells, where neither was found in Pdgfra^+^ (Fig. 7I; Suppl. 9G). *De novo* motif analysis similarly identified an OCT4:SOX17 motif uniquely in primed DP cells, alongside the early endoderm and TE related pluripotency factor, KLF5^38^ (Suppl. 9H), while active Oct4^+^ enhancers contain OCT4-SOX2-NANOG motifs alongside those that recognise downstream components of JAK/STAT (Suppl. 9I). Overall, our findings suggest a model where DP cells are maintained in a state primed for multi-lineage differentiation through persistence of a unique set of lineage-specific transcription factors, where absence of phosphorylated STAT3 leads to decommissioning of these regulatory regions results in commitment (Fig. 7J).

## Discussion

In this paper, we have demonstrated that the cells of the early PrE are sufficient to build an entire mouse and we provide novel molecular insight into pioneering work on the regulative properties of the mammalian preimplantation embryo^4^. As we found that an ICM composed entirely of Epi cells had a limited ability to reconstruct a blastocyst, our work suggests that the PrE may be central to this unique regenerative phenomenon. The capacity of the PrE to regenerate all cell types of the blastocyst is governed by the status of lineage specific enhancers in these cells and the persistence of key transcription factors, despite the onset of early differentiation. We find that OCT4 is key to this process as a factor that sits at the pinnacle of the gene regulatory network governing plasticity in addition to a well-known role in supporting pluripotency.

The amount of OCT4 expressed in reprogramming and self-renewal determines cell identity, where in ESCs, high and low levels of OCT4 enable differentiation towards endoderm and mesoderm or trophectoderm, respectively, and moderate levels promote pluripotency^37, 71^. In eutherian mammals, it is a single gene, *Pou5f1*, that supports this diversity of function by both activating and repressing transcription^72^. Given that these roles encompass multiple levels of differentiation, it is not surprising that *Oct4* is extensively conserved in evolution, where a number of species retain two paralogue POU domains, POU5F1 and POU5F3^73–76^. In somatic cell reprogramming, *Oct4* exhibits pioneer-like activity, where it recognizes sites in chromatin consistent with its ability to bind nucleosome assembled DNA^77^, as well as binding at enhancers associated with signalling-dependent differentiation genes, including those that respond to TGFβ and ERK^78, 79^. What is the ubiquitous role for *Oct4* in all these cell types? Perhaps as we describe here, *Oct4* functions as a neutral transcription factor, priming enhancers for lineage-specific response to signalling, suggesting that it is more of an enabler than a direct driver of transcription. Thus, in DP nEnd, OCT4 safeguards cell type-determining Epi, PrE and TE regulatory regions, preparing these for future activation.

If OCT4 is a passive player, then what factors mediate the conversion of DP nEnd into an Epi or TE-like cell type? We found that expansion of the DP population and subsequent PrE-to-Epi transition, both *in vitro* and *in vivo*, requires JAK/STAT signalling. As LIF signalling is associated with diapause where differentiation is indefinitely stalled^80^, the support of this signalling pathway within the DP population could reflect a general role in blocking commitment and expansion of a plastic cell type. The dependence on oxidative phosphorylation in DP nEnd is also consistent with the capacity of increased fatty acid oxidation to support embryo longevity and enhanced diapause via suppression of glycolysis^81^. As we observe here that the DP population proliferates faster than Pdgfra^+^ nEnd, this might explain our earlier observations that LIF signalling supports expansion of the PrE compartment *in vivo*^47^. We demonstrate that JAK/STAT signalling is required for the competence to undergo PrE-Epi conversions^9, 12, 13^, supporting an evolutionarily conserved role for JAK/STAT signalling in reprogramming, regeneration, proliferation and survival^51, 82, 83^, whereby a block to JAK attenuates plasticity by enabling commitment in the endoderm. Thus, while FGF/ERK signalling governs the Epi-PrE lineage decision during normal embryonic development and PrE differentiation, we find that JAK/STAT signalling is required to preserve plasticity and allow the reactivation of Epi and TE networks during aberrant development.

Plasticity in the ICM appears to be limited by the exposure to sustained FGF/ERK signalling^71^, and while it was previously suggested that the PrE becomes committed following FGF stimulation^32^, in this study, the duration of FGF4 treatment is shorter followed by a longer length of time for regeneration. While ESCs are generally considered closest to blastocyst-stage Epi, certain rare populations maintain the capacity to support multi-lineage competence for PrE and TE^34, 84^. Although not canonically defined as totipotent, as these cells cannot generate an embryo, we have referred to these as experimentally totipotent as they co-express certain PrE and Epi determinants that set up priming for multi-lineage differentiation^23^. The presence of a similar subpopulation in nEnd suggests that our *in vitro* culture traps developmental transition states that may be relatively fleeting *in vivo*. Similarly, this explains why we did not originally detect this heterogeneity in nEnd culture^28^, as we did not possess the appropriate reporter cell lines to purify subpopulations. Using an OCT4 reporter and appropriate cell surface markers, DP nEnd can be repeatedly isolated by FACS and introduced into different culture systems, allowing these cells to be used as a model along the spectrum between canonical pluripotency and totipotency.

What are the trajectories of cell state transitions during reprogramming and differentiation and how do these relate to Oct4? The emergence of a XEN-like intermediate during conversion of somatic cells to iPSCs, either en route to pluripotency^54, 85, 86^, or alternatively as a by-product of reprogramming^53^, has been widely reported. While these endodermal reprogramming intermediates were initially reminiscent of nEnd, our analysis of transcriptomic data demonstrates that these XEN-like intermediates resemble postimplantation-stage endoderm or XEN cells and that subpopulations of nEnd fall along the trajectory to acquisition of pluripotency. Moreover, our data suggests that the transition between DP and Oct4^+^ nEnd does not occur through reprogramming via a 2C-like intermediate, but rather as a direct conversion between lineages, in agreement with recent reports^45, 87^.

While the TE is an innovation of eutherian mammals, the PrE (also known as hypoblast) and its descendants originate deep in amniote evolution. The PrE has long been associated with yolk sac formation, anterior-posterior axis formation, and positioning of the starting point for gastrulation^88, 89^. In addition to these functions, why does the early PrE maintain a plastic population in preimplantation development? We suggest that the embryo needs a reservoir as a developmental insurance policy in case of aberrant development. Following the evolution of the placenta, the function of the yolk sac can be seen as somewhat redundant, but as we find its founder cells are required for regenerative purposes in preimplantation development, this could explain its conservation. Unlike the TE, where aneuploidy is common, the PrE is composed of normal cells that actively participate in elements of embryonic gut development^36, 38^. These cells therefore represent an ideal source of replacement cells to compensate for damage during preimplantation development.

## Acknowledgements

We thank N. Festuccia for EKOiE ESCs. We thank J. Martin Gonzalez, R. A. L. Barraza and the Core Facility for Transgenic Mice for embryo generation, training and support; H. Wollmann, M. Michaut and the reNEW Genomics Platform for technical expertise, support and use of instruments; G. de la Cruz, P. van Dieken and the reNEW Flow Cytometry Platform for training, technical expertise, support and use of instruments; E. F. Rebollo, M. Paulsen and the reNEW Cell Culture Platform for support and use of instruments. We also thank Jan J. Zylicz and members of the Brickman lab for fruitful discussions and critical comments on this manuscript. The work was funded by the Lundbeck Foundation (R370-2021-617, R198-2015-412 and R286-2018-1534), the Independent Research Fund Denmark (DFF-8020-00100B), the Danish National Research Foundation (DNRF116) and the Novo Nordisk Foundation (NNF17CC0027852 and NNF21CC0073729).

## Author contributions

Conceptualization, M.L.-A. and J.M.B.; Methodology, M.L.-A., A.C.S., A.R.-R., T.E.K., and J.M.B.; Investigation, M.L.-A., A.R.-R, A.C.S., T.E.K. and M.P.; Formal Analysis, M.L.-A., M.P. and T.E.K.; Data Curation, M.L.-A., M.P. and T.E.K.; Visualization, M.L-A. and M.P.; Writing – Original Draft, M.L.-A. and J.M.B.; Supervision, J.M.B.; Funding Acquisition, M.L.-A., M.P. and J.M.B.

## Methods

### Mouse maintenance and embryo collection

C57BL/6N mice were housed under 12h light/dark cycles at RT of 22°C (± 2°C) and a humidity of 55% (±10%), with the air in room changed 8-10 times per hour. Natural mating was set up in the evening and mice were checked for copulation plugs the following morning, which was determined as embryonic day 0.5 (E0.5). The males used for mating were 8-60 weeks old, while females were 8-30 weeks old. 8-cell embryos were collected by flushing the oviducts at E2.5. Zona pellucidae was removed from all embryos not used for chimera generation by incubation with acidic Tyrode’s solution (Sigma Aldrich) for 1-3min at RT. All animal work was carried out in accordance with European legislation. All work was authorized by the Danish National Animal Experiments Inspectorate (Dyreforsøgstilsynet, license no. 201815-0201-01520) and performed according to national guidelines.

### Blastocyst immunosurgery

Immunosurgery was performed on E3.5 blastocysts following removal of the zona pellucida. The embryos were placed in 20% anti-mouse rabbit serum (Sigma Aldrich) for 1h, after which they were transferred to 20% guinea pig complement (Sigma Aldrich)^31^. The lysed TE cells were then removed mechanically from the ICM by mouth pipetting. ICMs that disintegrated within 1-2h following immunosurgery were not included in final analysis.

### Immunostaining of E4.5 embryos

Embryos between E2.5-4.5 were fixed with 4% paraformaldehyde (PFA) for 10-15min at RT with gentle rocking and washed 3 times in PBS containing 5mg/mL polyvinylpyrrolidine (PBS/PVP). The embryos were then transferred to a Nunc MicroWell MiniTray (Thermo Scientific), permeabilized with permeabilization buffer (PBS/PVP and 0.25% Triton X-100) for 30min at RT and blocked with blocking buffer (PBS/PVP, 3% donkey serum, 0.1% BSA and 0.1% Triton X-100) for at least 15min at RT or overnight at 4°C in the dark. Primary antibodies were diluted in blocking buffer and incubated overnight at 4°C in the dark, followed by 3 washes with embryo blocking buffer for 10min each. Secondary antibodies were all diluted in embryo blocking buffer at a concentration of 1:200 and incubated for 1h at RT, followed by 3 washes with embryo blocking buffer for 10min each. Where applicable, DAPI was added at 1:5000.

### Immunostaining of E6.5 embryos

E6.5 embryos were timely dissected from the uterus in PBS at RT, fixed with 4% PFA for 30min at RT with gentle rocking and washed 3 times for 15min in PBS. Afterwards, embryos were permeabilised for 30min in permeabilization buffer (PBS/PVP and 0.5% Triton X-100) and blocked with blocking buffer (PBS/PVP with 2% donkey serum, 0.1% BSA and 0.1% Tween20). Primary antibodies were diluted in blocking buffer and incubated for 48h at 4°C in the dark, then washed 3 times for 15min and overnight at 4°C in the dark. Secondary antibodies were diluted in blocking buffer and incubated for 24h at 4°C in the dark, then washed 3 times for 15min and overnight at 4°C in the dark. DAPI was added to blocking buffer at 1:5000 and incubated for 3h at RT in the dark, then washed 3 times for 15min.

### Mouse ESC culture and differentiation to PrE

ESCs were maintained in naïve conditions on gelatinised tissue culture plates in N2B27 medium: Neurobasal (Gibco), DMEM/F-12 (Gibco), B-27 (Gibco), N2 (Gibco), L-glutamine (Gibco) and 2-mercaptoethanol (Sigma) supplemented with 2iLIF: 10ng/mL LIF (made in house), 3µM CHIR99021 (Axon Medchem) and 1µM PD0325901 (Sigma)^24^. For PrE differentiation, 25×10^3^ cells per/cm^2^ cells were seeded on gelatinised tissue culture plates for 24h in PrE basal medium: RPMI 1640 (Gibco) basal medium and B-27 minus insulin (Gibco). PrE basal medium was then supplemented with 10ng/mL LIF, 3µM CHIR99021 and 10ng/mL Activin A (Peprotech) (RACL medium) for an additional 4-6 days^28^. All cells were maintained under normoxic conditions at 37°C. Cell lines used in this study are listed in Suppl. Table 1.

### Expansion of nEnd

At day 5-7 of PrE differentiation, cells were collected at single-cell suspension and stained for PDGFRA-APC, after which PDGFRA-APC positive cells were isolated by FACS. E14 WT PrE cells were additionally stained for PECAM-FITC to exclude residual ESCs. GCSG PrE cells were sorted by expression of GATA6-mCherry. 15-20×10^3^ cells per/cm^2^ of PDGFRA APC or GATA6-mCherry positive cells were then seeded on MEF-coated tissue culture plates in RACL medium and passaged every 3-6 days. For 3D nEnd culture, AggreWell 400 plates were prepared as per the manufacturer’s instructions (STEMCELL Technologies). GATA6-mCherry^+^ or PDGFRA-APC^+^ nEnd isolated by FACS were then plated such that there were ∼50-100 cells per single AggreWell for a total of 6-12×10^4^ cells in each well containing RACL medium. Each well was then pipetted gently 2-3 times to allow even distribution of cells and the plate centrifuged at 100 x *g* for 3min. RACL medium was changed every 2 days. All cells were maintained under normoxic conditions at 37°C.

### Flow cytometry

Cells were dissociated to single-cell suspension with Accutase (STEMCELL Technologies), incubated with the appropriate conjugated antibody (Suppl. Table 1) for 25min in PBS supplemented with 6% FBS at 4°C on ice and washed several times. DAPI was added for live/dead discrimination and cells were either analysed on a LSRFortessa (BD Biosciences) or sorted on either an Aria III (BD Biosciences) or SH800Z (Sony). Data was analysed using FACSDiva (BD Biosciences) and FCS Express 7 (De Novo Software).

### Generation of chimeras

OCT4-mCherry-H2B-Venus DP and Pdgfra^+^ nEnd was isolated by FACS, after which 5 cells of either population was injected into 8-cell morulae (E2.5). Injected embryos were then cultured until E4.5 *in vitro* in KSOM under normoxic conditions at 37°C.

### Immunostaining of cells

Cells were washed with PBS and fixed with 4% PFA at 37°C for 10min, permeabilised with 1% Triton for 1h at RT and blocked with 3% donkey serum, 0.1% BSA and 0.2% Triton for 1h at RT or overnight at 4°C. Primary antibody incubation was done overnight at 4°C, followed by secondary antibody incubation for 1hr at RT and nuclear stain with DAPI.

### RNA preparation and RT-qPCR

Total RNA was extracted using either the RNeasy Mini Kit (Qiagen) or RNeasy Micro Kit (Qiagen) and cDNA was generated using SuperScript III reverse transcriptase (Invitrogen) and Random Hexamer primers (Invitrogen). RT-qPCR was performed using the LightCycler 480 Instrument II (Roche) with the primers and probe pairs listed in Suppl. Table 1 using the Universal Probe Library system. Relative concentrations were determined and normalised to the geometric mean of two housekeeping genes.

### Library preparation for scRNA-seq and transcriptome sequencing

2iLIF ESCs, nEnd and 3D nEnd were dissociated to single-cell suspension with Accutase (STEMCELL Technologies) and re-suspended in PBS supplemented with 6% FBS and 1:5000 DAPI for live/dead discrimination. DAPI negative cells were isolated by FACS for a total of 20×10^4^ cells per sample. Libraries were prepared and sequenced using 10X Genomics 3’ CellPlex Multiplexing solution. Samples were demultiplexed to GEX1 and GEX2 using cellranger (v6.1.12) mkfastq. For each sample, a cellranger multi was executed to retrieve the count matrices with mm10-2020-A as a reference genome. Configuration files are included in the GitHub link. Spliced, unspliced and ambiguous reads were determined from aligned BAM files from cellranger using STARSolo v2.7.9a for each sample.

### scRNA-seq pre-processing, filtering, and quality control

The raw dataset comprising 21,631 cells and 32,285 genes was converted to a Seurat (v4.3.0) object and filtered for cells with high mitochondrial content, low gene number and outliers, after which the dataset contained 14,788 cells. The raw UMI counts were normalized using ‘NormalizeData’ followed by ‘FindVariableFeatures’, which identified 2000 highly variable genes. The counts were further scaled using ‘ScaleData’ with default settings to perform PCA dimensionality reduction (‘Run PCA’). Shared nearest neighbour graph was computed using the first 20 principal components, followed by identifying 13 clusters with resolution 0.6 using Louvain clustering, visualised using UMAP.

### RNA velocity

STARSolo output matrices were merged with the processes count matrix based on common cell barcodes. RNA velocity analysis was performed using scVelo^39^ following the recommended workflow. We have adjusted n_neighbors and n_pcs to 20 and used 2000 highly variable genes with dynamical modelling to estimate cellular dynamics. The velocities were projected on already computed UMAP visualisation. To infer latent time that matched the experimental setup, we set cluster 7 (DP) as a starting point.

### Integration of scRNA-seq with published data

Based on an extensive benchmarking review on integration tools^90^, we integrated the Nowotschin 2019 dataset using SCVI^56^. We subset Nowotschin 2019 to E3.5 and E4.5 stages and merged them with our experiment. Raw counts were used with the following settings: dropout_rate = 0.2, dispersion = gene and gene_likehood = nb. We validated the integration by performing trajectory analysis using PAGA ^57^ on the embedding space.

### siRNA transfection

nEnd culture was sorted by FACS for PDGFRA-APC single positive cells and seeded at 15×10^3^ cells/cm^2^ in RACL on MEF-coated plates. Transfection with SMARTPool ON TARGETplus Pou5f1/Oct4 siRNA (Horizon Discovery) and ON-TARGETplus Non-targeting Control siRNAs Control #1 (Horizon Discovery) was performed using Lipofectamine RNAi MAX (Life Technologies) at a final concentration of 100nM. siRNAs were added 48h after seeding and collected for analysis after an additional 48h.

### CUT&Tag

CUT&Tag was performed as previously described^63, 91^ with slight alterations. All samples were isolated by FACS based on OCT4-mCherry and PDGFRA-APC expression, and 25×10^4^ cells per sample replicate was used for downstream processing. Primary antibody incubation was done at 4°C overnight, Tn5 was purchased from EMBL Heidelberg and DNA precipitation was performed over the weekend at -80°C. DNA was amplified with 14 PCR cycles and the samples were sequenced paired end on an Illumina NextSeq 500. Downstream analysis was performed using Bedtools to determine BedGraph coverage, peaks were then called against an IgG control using SEACR^92^ using the ‘stringent’ parameter, and consensus peaks were determined used DiffBind^93^, visualized with Deeptools^94^. BAM files were merged for visualisation using SAMtools^95^ and annotation of peaks was performed using HOMER^96^.

### Microscopy and imaging analysis

Cells were imaged using either a confocal Leica TCS SP8, confocal Leica STELLARIS or widefield Leica AF6000 microscopes. Image analysis was carried out using FIJI (ImageJ), Imaris v9.9.1 (Oxford Instruments) and CellProfiler^97^. All image quantification was performed on single optical sections of original images from multiple images taken across different biological replicates. Maximum projections were used only for visualisation purposes.

### Statistical analysis and reproducibility

All embryo treatments were performed using minimum two independent litters at different times, where control embryos were littermates to the corresponding treatment for a given experiment. Statistical tests were performed using R and GraphPad Prism 9. Unless otherwise stated, experiments were performed with three biological replicates (*n* = 3), *P* values were determined by standard *t-*test or one-way ANOVA, where appropriate, and error bars indicate ± standard deviation. Significance was defined as: not significant (ns) ≥ 0.05; \**P* < 0.05; \*\**P* < 0.01; \*\*\**P* < 0.001.

## Supplemental information

**Supplementary Figure 1: Quantification of control and treatment conditions, related to Figure 1**

**(A)** Table summarising *n* values and success rates for treatments performed in Fig. 1A-F; S1E. **(B)** Representative immunostaining of an E3.5 embryo treated with FGF4 for 24h for indicated markers including DAPI for visualisation of nuclei, imaged by confocal microscopy. White arrowheads indicate cells expressing GATA6 and low levels of NANOG. **(C)** Representative immunostaining of E3.5 embryos based on Fig. 1A for indicated markers including DAPI for visualisation of nuclei, imaged by confocal microscopy. **(D)** Single cell quantification of embryos in **C** for NANOG and GATA6 immunostaining normalized to DAPI. *n* values indicate total number of embryos quantified. **(E)** Representative immunostaining of an embryo 48h following immunosurgery for CDX2 and DAPI for visualisation of nuclei, imaged by confocal microscopy. White arrowheads indicate morphologically stressed cells expressing CDX2. **(F)** Lineage allocation of E4.5 embryos following immunosurgery in control and FGF4-treated embryos. **(G)** Table summarising *n* values and success rates for treatments performed in Fig. 1G-J. **(H)** Representative photograph of pups derived from E3.5 PrE detailed in Fig. 1G. **(I)** Schematic of workflow to establish viability of embryos at E6.5 following trophectoderm reconstruction in FGF4-treated embryos compared to control. **(J)** Table summarising *n* values and success rates for treatments performed in Fig. 1K, M; S1I. **(K)** Brightfield images of uterine horns (left), deciduae (middle) and embryos (right) for control transfers and FGF4-treated embryos. White arrowheads indicate position of deciduae within uterine horn. Scale bars represent 50µm.

**Supplementary Figure 2: PrE and nEnd differentiation competence of GCSG reporter cells and derivation from E3.5 ICMs, related to Figure 2**

**(A)** Brightfield and immunofluorescence imaging of GCSG reporter cells in 2iLIF and day 7 of PrE differentiation, imaged by confocal microscopy. **(B)** Flow cytometry contour plots of **A** where bottom left quadrant indicates gating based on a negative control. **(C)** Representative flow cytometry contour plots of GCSG nEnd showing gating strategy for isolating GATA6 mCherry^+^ and SOX2-GFP^+^ populations for re-plating. **(D)** Representative flow cytometry contour plots of E14JU nEnd stained for PDGFRA-APC and PECAM-FITC followed by FACS for PDGFRA-APC^+^ cells and analysed by flow cytometry after 96h. Bottom left quadrant indicates gating based on a negative control. **(E)** Quantification of 3D nEnd growth in size 24-144h following FACS and seeding in AggreWell’s (*n* = 200 per timepoint). **(F)** Quantification of total expression by flow cytometry of GCSG 3D nEnd 0-120h following FACS for GATA6-mCherry^+^/SOX2-GFP^-^ cells. **(G)** Representative brightfield images of nEnd derivation from E3.5 blastocysts. **(H)** Representative flow cytometry contour plots of embryo derived nEnd stained for PDGFRA-APC and PECAM-FITC. Bottom left quadrant indicates gating based on a negative control. **(I)** Representative immunostaining of 3D nEnd from embryo-derived nEnd for indicated markers including DAPI for visualisation of nuclei, imaged by confocal microscopy. Scale bars represent 50µm in **A**, **I**; 100µm in **G**.

**Supplementary 3: OCT4 expression separates uncommitted from committed extra embryonic endoderm cell types, related to Figure 3**

**(A)** t-distributed **stochastic** neighbour (tSNE) embedding of scRNA-seq of the mouse preimplantation embryo at E3.5-4.5^38^, where colour scale represents expression of Pou5f1 transcripts and shape indicates cell type. **(B)** Immunostaining of E14JU nEnd for indicated markers including DAPI for visualisation of nuclei, imaged by confocal microscopy. **(C)** Flow cytometry contour plot of unstained OCT4-mCherry nEnd where the bottom left quadrant indicates gating based on a negative control. **(D)** Flow cytometry contour plot of OCT4 mCherry nEnd stained for PECAM-FITC and PDGFRA-APC, where gating of PECAM-FITC^+^ cells (left) are visualised for OCT4-mCherry and PECAM-FITC co-expression (right). **(E)** Representative flow cytometry contour plots for monitoring OCT4-mCherry, PECAM-FITC and PDGFRA-APC expression from 24-96h following FACS for PDGFRA-APC^+^ cells. Bottom left quadrant indicates gating based on a negative control. **(F)** Representative brightfield imaging and immunostaining of OCT4-mCherry nEnd for E-cadherin and DAPI for visualisation of nuclei, imaged by confocal microscopy. **(G)** Cell size of Pdgfra^+^ (*n* = 333) and DP (*n* = 511) nEnd based on E-cadherin immunostaining. **(H)** Nuclear size of Pdgfra^+^ (*n* = 426) and DP (*n* = 400) nEnd based on DAPI localisation. **(I)** Quantification of doubling time in Pdgfra^+^ and DP nEnd based on total cell number after 72h in culture. **(J)** Immunostaining of OCT4-mCherry nEnd for indicated markers including DAPI for visualisation of nuclei, imaged by confocal microscopy. Scale bars represent 50µm.

**Supplementary 4: XEN cells represent later-stage extra-embryonic endoderm lacking OCT4 expression that cannot be rescued upon culture in nEnd medium**

**(A)** Schematic of E14JU ESCs in 2iLIF towards XEN cells^26, 43^ followed by FACS for PDGFRA-APC^+^ cells to purify endoderm population. **(B)** Flow cytometry contour plots of XEN cell derivation compared to 2iLIF ESCs and nEnd stained for PECAM-FITC and PDGFRA-APC. Bottom left quadrant indicates gating based on a negative control. **(C)** Brightfield images of XEN derivation compared to 2iLIF ESCs and nEnd. **(D)** Immunostaining and brightfield images of nEnd and XEN cells for indicated markers including DAPI for visualisation of nuclei, imaged by confocal microscopy. **(E)** Flow cytometry contour plots of XEN cells cultured in indicated conditions stained for PECAM-FITC and PDGFRA-APC expression. Bottom left quadrant indicates gating based on a negative control. **(F)** Immunostaining of cXEN cells for indicated markers including DAPI for visualisation of nuclei, imaged by confocal microscopy. **(G)** RT-qPCR of XEN cells compared to nEnd isolated by FACS for PDGFRA-APC expression for indicated markers. Scale bars represent 100µm in **C**; 50µm in **D**, **F**.

**Supplementary 5: nEnd differentiates into a TE-like cell type**

**(A)** Brightfield images of nEnd and nEnd cultured in TSC medium. **(B)** Immunostaining of OCT4-mCherry nEnd and nEnd cultured in TSC medium for indicated markers including DAPI for visualisation of nuclei, imaged by confocal microscopy. White arrowheads point to BRACHYURY/CDX2 co-expressing cells and white asterisk points to CDX2 single positive cells. **(C)** Schematic showing treatment of nEnd in either RACL or TSC medium followed by whole transcriptome analysis by scRNA-seq. **(D)** UMAP dimensional embedding of 11,194 nEnd cells in either RACL or TSC medium. Top: colouring based on culture condition defined in **C**; bottom: colouring based on Louvain clustering. **(E)** Allocation of each culture condition per cluster. **(F)** UMAP dimensional embedding showing single cell expression of indicated markers. **(G)** Heatmap of candidate lineage markers expressed in log2 normalized clustered data. Scaled by row. **(H)** Expression of indicated TE and Mesoderm-specific markers in TE-like (*n* = 357 cells) and Mes-like (*n* = 221 cells) clusters. **(I)** Differential expression of TE and mesoderm candidate markers in TE-like and Mes-like clusters. **(J)** Volcano plot of DEGs between Mes-like and TE-like clusters representative of log2 fold change > 0.25 and *p* < 0.05. **(K)** Comparative gene expression overlap analysis of TE-like and Mes-like clusters with scRNA-seq of the mouse preimplantation embryo^38^. Grey points represent *p* > 0.05. Scale bars represent 100µm in A; 50µm in B.

**Supplementary 6: Loss of JAK/STAT signalling results in the loss of the DP population, related to Figure 4**

**(A)** Brightfield images of OCT4-mCherry nEnd in control and treated conditions. Arrowheads indicate aggregates containing putative reverted cells and white dashed line highlights epithelial nEnd with compact morphology. **(B)** Single cell quantification of embryos for OCT4 and pSTAT3 immunostaining outlined in Fig. 4A normalized to DAPI. *n* values indicate total number of cells quantified. **(C)** Representative flow cytometry contour plots of OCT4 mCherry nEnd stained for PDGFRA-APC showing gating strategy for isolating either PDGFRA-APC^+^ or OCT4-mCherry^+^ cells for downstream analysis. Bottom quadrant indicates gating based on negative control. **(D)** Representative flow cytometry contour plots of PDGFRA-APC^+^ OCT4-mCherry nEnd in control and treated conditions using gating strategy in **C**. Bottom quadrant indicates gating based on negative control. Scale bar represents 100µm.

**Supplementary 7: Clustering annotation strategy for 2iLIF, nEnd and 3D nEnd scRNA seq, related to Figure 5**

**(A)** Schematic showing scRNA-seq strategy for nEnd and 3D nEnd. **(B)** Allocation of each culture condition per cluster. **(C)** Differential expression of candidate 2C markers across all clusters. **(D)** Heatmap showing expression of genes related to apoptosis^98^ in log_2_ normalized clustered data. **(E)** Gene ontology analysis of selected biological processes of upregulated genes in the putative Apoptotic cluster. **(F)** PCA of scRNA-seq dataset for 2iLIF, nEnd and 3D nEnd. Left: colouring based on culture condition; right: colouring based on Louvain clustering. **(G)** Heatmap of candidate lineage markers in log_2_ normalized clustered data. Scaled by row. **(H)** Heatmap of scaled expression of candidate early (Pou5f1, Tbx3, Fgfr2, Klf5, Idh1, Gata6) and late (Gata4, Col4a1, Hnf4a, Foxa2, Vegfa, Cited1) PrE genes from scRNA-seq of ICM to PrE *in vivo* across pseudotime^38^. Scaled by row. **(I)** PAGA of integrated *in vitro* nEnd, 3D nEnd and nEnd to TSC (this study), and somatic cell reprogramming to iPSCs from a XEN-like intermediate^55^ datasets using SCVI^56^. Colouring based on Louvain clustering (XEN-like 1-3, Pdgfra^+^ 1-5, DP, Oct4^+^, Mes-like, TE-like) or reprogramming stage (XEN, SII D8, SII D12, SIII D3, SIII D6, SIII D8, SIII D10, SIII D15, SIII D21). Thicker lines indicate highly connected regions and thinner lines indicate regions with lower confidence.

**Supplemental 8: Gene expression and trajectory inference in 2iLIF, nEnd and 3D nEnd scRNA-seq, related to Figure 5**

**(A)** Volcano plot of DEGs between DP and Pdgfra^+^ clusters representative of log_2_ fold change > 0.25 and *p* < 0.05. **(B)** Representative flow cytometry contour plots gated for PDGFRA-APC expression of OCT4-mCherry nEnd cultured in 2DG or Etomoxir for 5 days. **(C)** Quantification of PDGFRA-APC expression in nEnd treated with 2DG or Etomoxir compared to control. **(D)** Euler diagram for overlap of DEGs in Oct4^+^, DP and Pdgfra^+^ 1-5 clusters with 700 downstream targets of Oct4^59^. Candidate shared genes between DP and Oct4^+^ highlighted. **(E)** Volcano plot of DEGs between Oct4^+^ and DP clusters representative of log_2_ fold change > 0.25 and *p* < 0.05. **(F)** UMAP dimensional embedding of nEnd and 3D nEnd cells by scRNA-seq, where clustering corresponds to Fig. 5A. **(G)** UMAP dimensional embedding of **F** where colouring represents RNA-velocities determined by scVelo and superimposed arrows represent inferred directionality of the dataset.

**Supplemental 9: Chromatin states in nEnd subpopulations demonstrate unique developmental characteristics, related to Figure 7**.

**(A)** Profiles of H3K4me1 and H3K27ac for Oct4^+^, DP and Pdgfra^+^ nEnd upstream Nanog compared to defined pluripotency (yellow) enhancers^34^, highlighted in grey with -5 and -45 Nanog SE annotated in blue. Bigwigs generated from 3 biological replicates. **(B)** Profiles of H3K4me1 and H3K27ac for Oct4^+^, DP and Pdgfra^+^ nEnd at *Col4a1* and *Col4a2* loci compared to defined pluripotency (yellow) and nEnd (magenta) enhancers ^34^. Bigwigs generated from 3 biological replicates. **(C)** Position of active Oct4^+^ peaks annotated to the nearest TSS (±20kb) relative to genes selected for Fig. 7G and other downstream targets of Oct4^59^. **(D)** Position of primed DP peaks annotated to the nearest TSS (±20kb) relative to selected genes related to trophectodermal fate. **(E)** Euler diagrams showing overlap between annotated genes within ±20kb of primed DP peaks and DEGs in scRNA-seq for Oct4^+^, Pdgfra^+^ and TE-like clusters to DP nEnd. **(F)** Gene ontology analysis of selected biological processes for primed Pdgfra^+^, DP and Oct4^+^ peaks. **(G)** Ternary plot of motif enrichment across Oct4^+^, DP and Pdgfra^+^ nEnd in the repressed category. *p* < 0.05 for at least one population in every motif shown, where the axis represents -log10(*p*-value). Grey dots indicate all motifs and coloured dots highlight motifs of interest. **(H, I)** *De novo* sequence motifs for **H** primed DP and **I** active Oct4^+^ cells.

**Supplementary Table 1:** Details of cell lines, antibodies and primers used

