## Supplementary Figure 1 for "Enhancer status in the primitive endoderm supports unrestricted lineage plasticity in regulative development"

A

| Untreated | | Treated | | | | |
|---|---|---|---|---|---|---|
| A | B | C | D | E | F | G |
| 11 | 13 | 34/45 (75.6%) | 13/16 (76.9%) | 14/22 (63.6%) | 9/11 (81.2%) | 6/24 (25%) |

B

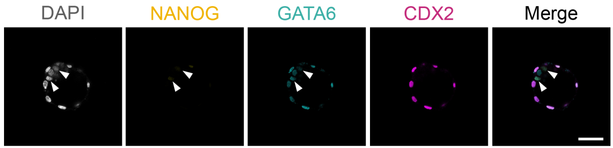


C

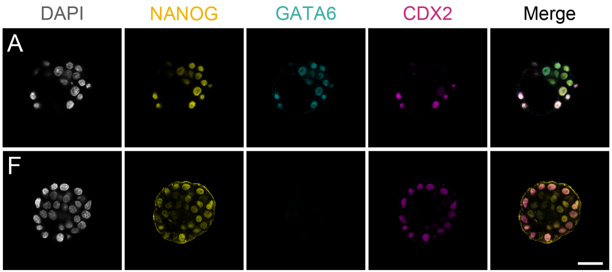


E

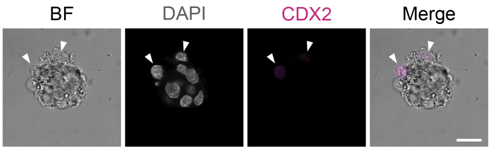


G

| Untreated | | Treated | | |
|---|---|---|---|---|
| A | B | H | I | J |
| 12 | 15 | 13/16 (81%) | 12/15 (80%) | 24/27 (89%) |

D

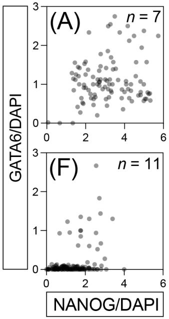


F

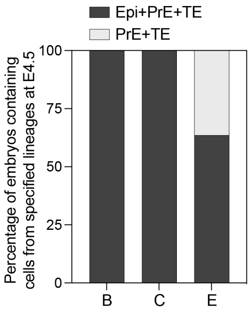


H

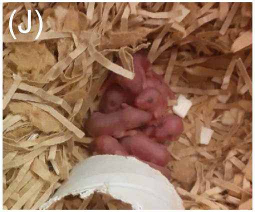


J

| | No implantation | Empty decidua | Embryo |
|---|---|---|---|
| Control transfer | 7 | 5 | 8 |
| -FGF4 -TE | 8 | 1 | 0 |
| +FGF4 -TE | 1 | 6 | 1 |

K

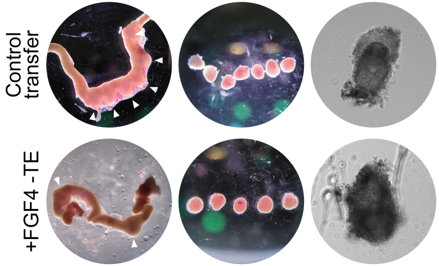


I

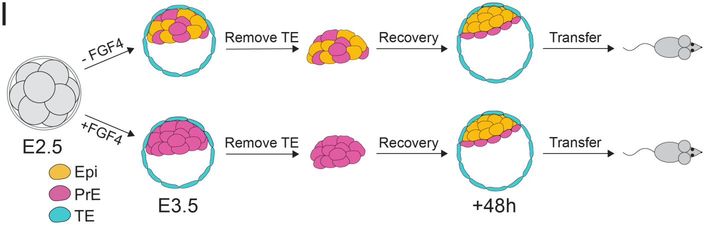
